# Allelic variants confer Arabidopsis adaptation to small regional environmental differences

**DOI:** 10.1101/2023.09.18.558200

**Authors:** Raúl Y. Wijfjes, René Boesten, Frank F. M. Becker, Tom P. J. M. Theeuwen, Basten L. Snoek, Maria Mastoraki, Jelle J. Verheijen, Nuri Güvencli, Lissy-Anne M. Denkers, Maarten Koornneef, Fred A. van Eeuwijk, Sandra Smit, Dick de Ridder, Mark G.M. Aarts

## Abstract

Natural populations of *Arabidopsis thaliana* provide powerful systems to study adaptation of wild plant species. Previous research has predominantly focused on global populations or accessions collected from regions with diverse climates. However, little is known about the genetics underlying adaptation in regions with mild environmental clines. We have examined a diversity panel consisting of 192 *A. thaliana* accessions collected from the Netherlands, a region with limited climatic variation. Despite the relatively uniform climate, we identified compelling evidence of local adaptation within this population. Notably, semidwarf accessions, due to mutation of the *GIBBERELLIC ACID REQUIRING 5* (*GA5*) gene, occur at a relatively high frequency near the coast and these displayed enhanced tolerance to high wind velocities. Additionally, we evaluated the performance of the population under iron deficiency conditions and found that allelic variation in the *FE SUPEROXIDE DISMUTASE 3* (*FSD3*) gene affects tolerance to low iron levels. Moreover, we explored patterns of local adaptation to environmental clines in temperature and precipitation, observing that allelic variation at *LA RELATED PROTEIN 1C* (*LARP1c*) likely affects drought tolerance. Not only is the genetic variation observed in a diversity panel of *A. thaliana* collected in a region with mild environmental clines comparable to that in collections sampled over larger geographic ranges, it is also sufficiently rich to elucidate the genetic and environmental factors underlying natural plant adaptation.

## Introduction

Adaptation is defined as the process through which a population attains higher fitness in its native environment than non-native populations sampled from different sites (Kawecki and Ebert, 2004). Unraveling the genomic and physiological basis of adaptation in plants is a central question in modern plant biology. From an evolutionary perspective, linking plant genotypes to adaptive traits helps to clarify how environmental gradients drive natural selection of both standing and novel variation, which ultimately may result in speciation (Sobel et al., 2010). It also enables us to model whether natural populations are able to adapt to future environmental conditions (Bay et al., 2017; Exposito-Alonso et al., 2019) and provides insights on processes or even genes to focus on when breeding crops for enhanced abiotic stress tolerance (Huang and Han, 2014).

The model plant species *Arabidopsis thaliana* provides an excellent system to study natural plant adaptation (Weigel and Nordborg, 2015). Recent studies of this species have shifted from studying the global HapMap population (Li et al., 2010), traditionally used for genome-wide association studies (GWAS), to populations sampled from regions containing a range of contrasting climates, including Sweden (Long et al., 2013), the Iberian Peninsula (Tabas-Madrid et al., 2018), and the south-west of France (Frachon et al., 2018), as these local populations are expected to minimize genetic heterogeneity and thus provide more statistical power for detecting quantitative trait loci (QTLs) (Brachi et al., 2013; Korte and Farlow, 2013). While studies on such regional populations have successfully identified several adaptive loci (Frachon et al., 2018; Tabas-Madrid et al., 2018; Arteaga et al., 2022), we know comparatively little about the loci driving adaptation in regions having a milder diversity in environmental conditions.

The Netherlands, a region which has remained largely under-sampled so far, provides an excellent opportunity to address this issue. This region covers an area comparable in size to the south-west of France, but much smaller than Sweden and the Iberian Peninsula. Compared to the other three regions, the Netherlands has only mild climatic clines (https://www.knmi.nl/klimaat-viewer), but past work indicated that local adaptation of *A. thaliana* may still be expected (Barboza et al., 2013). The Netherlands contains a moderate frequency of accessions with short inflorescences, mediated by the same loss-of-function allele of the *GA5* gene that managed to spread over more than 100 kilometres in the western part of the country (Barboza et al., 2013). There is reason to believe that more such signatures of local adaptation can be found, for instance based on the small differences in average temperature and annual precipitation, or the soil type on which *A. thaliana* grows in the Netherlands (De Vries et al., 2003). While it can be found on the clay and peat regions in the west and north of the country, it mainly grows on sandy soils, which largely occur in the east and south, and in the coastal dunes. It is often found in roadsides and gardens in which substantial movement of soils and seeds from other regions has occurred in the past.

To comprehensively address the contribution of genetic variation to plant adaptation in the Netherlands, we generated the <u>D</u>utch *<u>Ar</u>abidopsis <u>t</u>haliana* Map (DartMap) panel, a collection of 192 *A. thaliana* accessions sampled from different sites in the country. We show that this panel contains a surprisingly high level of genetic diversity and provide evidence of local adaptation to mild climatic clines. Moreover, we present strong evidence that variation at two loci is involved in mediating adaptation to wind tolerance and iron deficiency. Our work demonstrates that plant populations sampled in small geographic regions with a low diversity in environmental conditions can contain a level of genetic variation that is sufficiently rich to facilitate local adaptation.

## Results

### The degree of *A. thaliana* genetic diversity in the Netherlands is typical of that of most European regions

The global *A. thaliana* collection of the 1,001 Genomes (1001G) Project (The 1001 Genomes Consortium, 2016) includes only 11 lines from the Netherlands, which likely represents only a small portion of the region’s genetic variation due to limited dispersal of minor alleles (Tabas-Madrid et al., 2018). Therefore, we assembled and analysed the DartMap panel and used it together with the Dutch 1001G accessions to characterize the overall degree of genetic diversity of *A. thaliana* in the Netherlands. In total, we identified 2,712,612 SNPs and 353,974 indels in the DartMap panel and the Dutch 1001G accessions (collectively referred to as DartMap + 1001G panel for brevity) relative to the *A. thaliana* Col-0 nuclear genome reference sequence. Moreover, we detected 487 SNPs and 209 indels in the chloroplast sequence, and 137 SNPs and 15 indels in the mitochondrial sequence.

Furthermore, we examined copy number variation in the DartMap panel, excluding the Dutch 1001G accessions because they were sequenced on a different sequencing platform. We detected 29,155 copy number variants (CNVs) relative to the Col-0 reference genome. Most of these are deletions, less than 500 bp (Figures S1a and S1b), which is in line with the distribution of structural variants found in an earlier study on the full 1001G collection (Göktay et al., 2021). We found that 448 genes overlap with CNVs predicted to disrupt gene function and present at moderate frequencies within the DartMap panel (Table S1). We only considered deletions here, as only those were called with high precision (Figure S2). The 448 genes are significantly enriched for genes involved in disease resistance (*P* = 4.39×10^-6^).

To contextualize the degree of genetic diversity of the DartMap + 1001G panel, we compared it to similarly sized collections from Sweden (Long et al., 2013) and the Iberian peninsula (Tabas-Madrid et al., 2018), using the same SNP and indel calling approach. Considering only SNPs and indels that are polymorphic in the DartMap + 1001G panel (i.e. at least one accession contains a reference allele), the Swedish collection contains a similar number of variants, while the Iberian one is more diverse (Table 1). This disparity can be attributed to admixture of “relict” accessions in the Iberian collection, which originated from *A. thaliana* populations in Europe before the last ice age, and the genetically distinct “non-relict” group that repopulated Europe thereafter (The 1001 Genomes Consortium, 2016).

**Table 1:** The number of polymorphic bi-allelic SNPs and indels (“Variants”) found in regional *A. thaliana* populations.

| Population | Samples | Variants | Reference |
| --- | --- | --- | --- |
| Dutch (DartMap + 1001G) | 203 | 3,057,293 | This study, (The 1001 Genomes Consortium, 2016) |
| Sweden | 243 | 3,084,964 | (Long et al., 2013) |
| Iberian Peninsula | 190 | 4,482,982 | (Tabas-Madrid et al., 2018) |

Most SNPs and indels detected in the DartMap + 1001G panel are shared among multiple accessions and the distribution of their minor allele frequencies is similar to that of the Swedish collection (Figure 1a). In contrast, the Iberian collection contains a relatively larger fraction of rare variants (Figure 1a), possibly because the mountain systems in this region strongly limit the spread of minor alleles (Tabas-Madrid et al., 2018).

**Figure 1:**
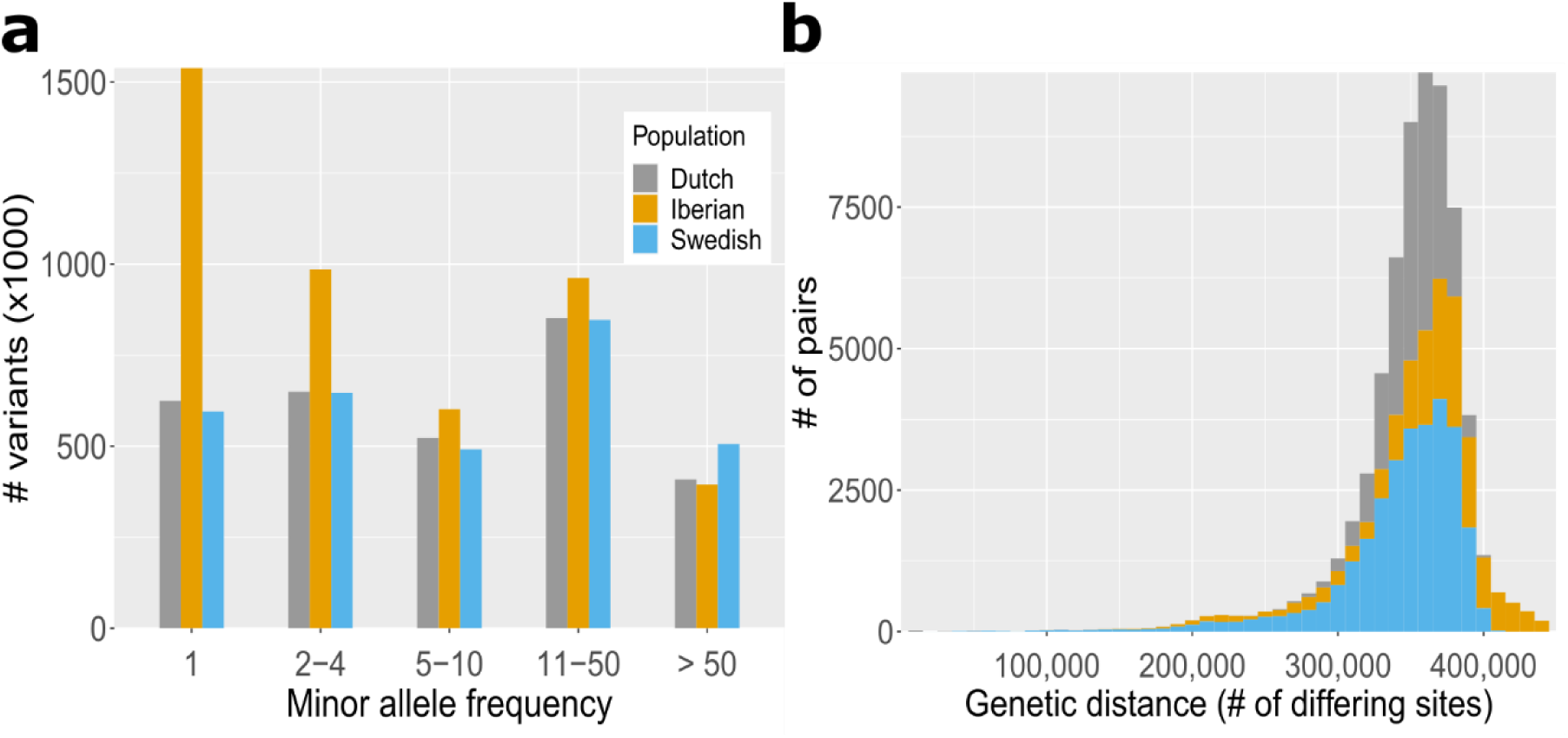
Comparing the genetic diversity of three regional populations. (a) Minor allele frequency of bi-allelic SNPs and indels. (b) Genome-wide similarity between pairs of samples (bars are overlapping).

To assess the genetic diversity of the DartMap + 1001G panel, we calculated pairwise genetic distances between individual accessions. Most pairs of accessions differ at approximately 320,000-340,000 variant sites, similar to the Swedish collection and most pairs of the Iberian collection (Figure 1b). However, the Iberian collection shows a right-skewed distribution of pairwise genetic distances, indicating the presence of accessions that are genetically more distant from each other compared to the majority (Figure 1b). This tail in the distribution aligns with previous findings of genetic distance between Iberian accessions and attributed to the presence of the relict accessions in this region (The 1001 Genomes Consortium, 2016). Taken together, our analyses indicate that genomic variation of *A. thaliana* in the Netherlands predominantly originated from the post-glacial expansion event from which most *A. thaliana* accessions in Europe descended (Lee et al., 2017).

### Genetic diversity of the *A. thaliana* in the Netherlands is shaped by ancient and contemporary forces

We explored the demographic and evolutionary history of the DartMap + 1001G panel using population structure analyses based on the genetic distances between individual accessions, which have proven useful for this purpose in previous studies of *A. thaliana* (The 1001 Genomes Consortium, 2016; Tabas-Madrid et al., 2018). We computed population structure of the DartMap + 1001G panel through hierarchical clustering based on pairwise organellar (Figure 2a, Table S2) and nuclear (Figure 2b, Table S2) genome-wide similarity, including all accessions of the 1001G collection sampled in Belgium, France, Germany, and the United Kingdom to evaluate relatedness between Dutch *A. thaliana* accessions and those of nearby countries. The dendrograms based on chloroplast and mitochondrial variation consist of three nearly equidistant groups (Figure S3) with identical spatial distributions across the Netherlands (Figure 2a), as expected for organellar sequences that are both predominantly maternally inherited. A previous study showed that European accessions contain several highly diverged chloroplast haplogroups of which the estimated time of divergence considerably predates the post-glacial expansion event (Hsu et al., 2019). A principal component analysis, based on chloroplast variation between individual accessions of the DartMap + 1001G panel and of a global collection of *A. thaliana*, separates Dutch accessions in three different groups (Figure 2c) that correspond to different subsets of chloroplast haplogroups identified in the same study (Figure 2d). This implies that the distinct organellar groups of the DartMap + 1001G panel represent ancient variation, while the nuclear groups reflect more recent genomic differentiation, explaining the limited overlap in spatial distribution between the two (Figure 2a and b).

**Figure 2:**
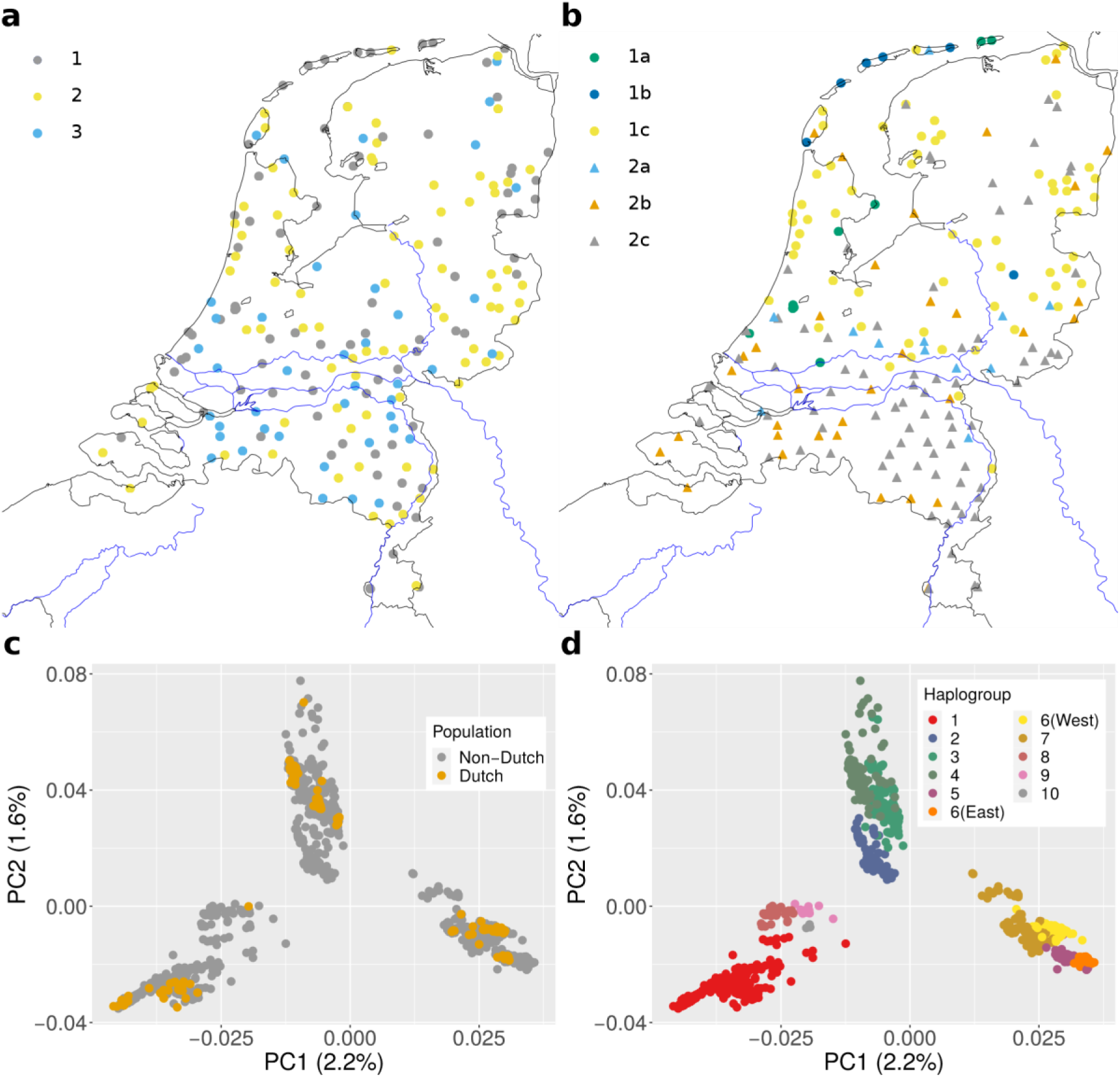
Population structure of *A. thaliana* in the Netherlands. (a-b) Geographical location of Dutch accessions clustered based on organellar (a) and nuclear (b) genome-wide similarity. Colours are used to distinguish accessions of different clusters. The main rivers of the Netherlands are depicted by blue lines. (c) Principal component analysis of chloroplast variants in Dutch accessions and the global collection of *A. thaliana* analysed in a previous study (Hsu et al., 2019). (d) Samples in (c) coloured according to the chloroplast haplogroups reported in a previous study (Hsu et al., 2019).

The spatial distribution of the nuclear groups provides insights into the factors that shaped genomic variation in the DartMap + 1001G panel following the postglacial expansion event. Based on nuclear variation, the panel can be divided into two main groups, predominantly separated by the Rhine and Meuse river branches (group 1 to the north and group 2 to the south) (Figure 2b). The rivers likely acted as natural dispersion barriers (Pico et al., 2008), restricting genetic flow between the two groups. It is worth noting that *A. thaliana* is generally not found on clay soil accompanying the rivers, except on sandy river dunes and residential areas where it has been introduced through human-induced transport of soil from other regions. Group 1 can be further divided into three subgroups (Figure 2b), of which two contain accessions that are closely related to each other (subgroups 1a and 1b) and less related to the rest of the population (Figure S4).

Subgroup 1a contains eight accessions of the Dartmap panel with short inflorescences, referred to as “semidwarfs”, and one accession (Tha-1) of the 1001G panel, previously described as a semidwarf (Barboza et al., 2013). The accessions from subgroup 1a are mainly found in the western part of the Netherlands, close to the North Sea coast (Figure 2b). Subgroup 1b is predominantly found on the Dutch islands in the north, implying that these islands were colonized by a limited number of founder accessions, mostly from group 1. Group 2 consists of accessions that are genetically similar to accessions from Germany (subgroup 2a), respectively France and the United Kingdom (subgroup 2b) (Figure S5), suggesting that these are part of a large genetic group spread over Northwestern Europe.

Besides large clusters, we identified 27 pairs of accessions that are near duplicates in terms of nuclear similarity (Table S3), with differences of fewer than 50,000 variant sites between them. This level of similarity is significantly lower than the median pairwise similarity of 354,388 variant sites (Figure 1a). Strikingly, they include 6 out of 8 of the semidwarf accessions of subgroup 1a, the furthest of which are separated by more than 100 km (Table S3) (Flood et al., 2016a).

### Semidwarf *ga5* mutants display tolerance to windy conditions

The widespread geographical distribution of highly identical genotypes that all have the same semidwarf phenotype suggests there may be a selective advantage to this trait, similar to the near-identical and widespread atrazine-resistant genotypes distributed along the United Kingdom railway system, which were identified previously (Flood et al., 2016a).

An earlier study already indicated a higher frequency of semidwarfs in the Netherlands compared to the global estimated frequency of at least 1% (Barboza et al., 2013). Next to the eight semidwarfs found in subgroup 1a, we identified a ninth semidwarf accession, Snh-1, in subgroup 2b. Almost all examined semidwarf accessions found in the field are loss-of-function mutants at the *GA5* locus, which encodes the *GIBBERELLIN 20-OXIDASE 1* (*GA20OX1*) gene (Barboza et al., 2013). Further investigation of the genomic sequences of *GA5* revealed that the eight semidwarf accessions in subgroup 1a, that are genotypically nearly identical, share a previously described *ga5* splice site mutation specific to Dutch semidwarfs (Barboza et al., 2013), while Snh-1 contains a 52-bp deletion in the last exon of *GA5*.

The appearance of a relatively high frequency of natural *ga5* mutants suggests that semidwarfism provides an adaptive advantage. Dwarfism in plants is a common adaptation to high altitude (Billings and Mooney, 1968; Grace, 1988; Luo et al., 2015), although it remains unclear which aspect of high altitude causes this. The majority of the Dutch semidwarfs are collected within 15 km of the coast, often only a few meters above sea level, so there is no association with high altitude. However, the locations where the semidwarfs were collected share an important characteristic providing a potential selective force, namely the presence of high wind speeds (Figure 3a). High wind speeds are also a characteristic of high-altitude environments. We therefore investigated whether the presence of high wind speeds could create a selective environment favouring semidwarf *A. thaliana* over normal-sized plants.

**Figure 3:**
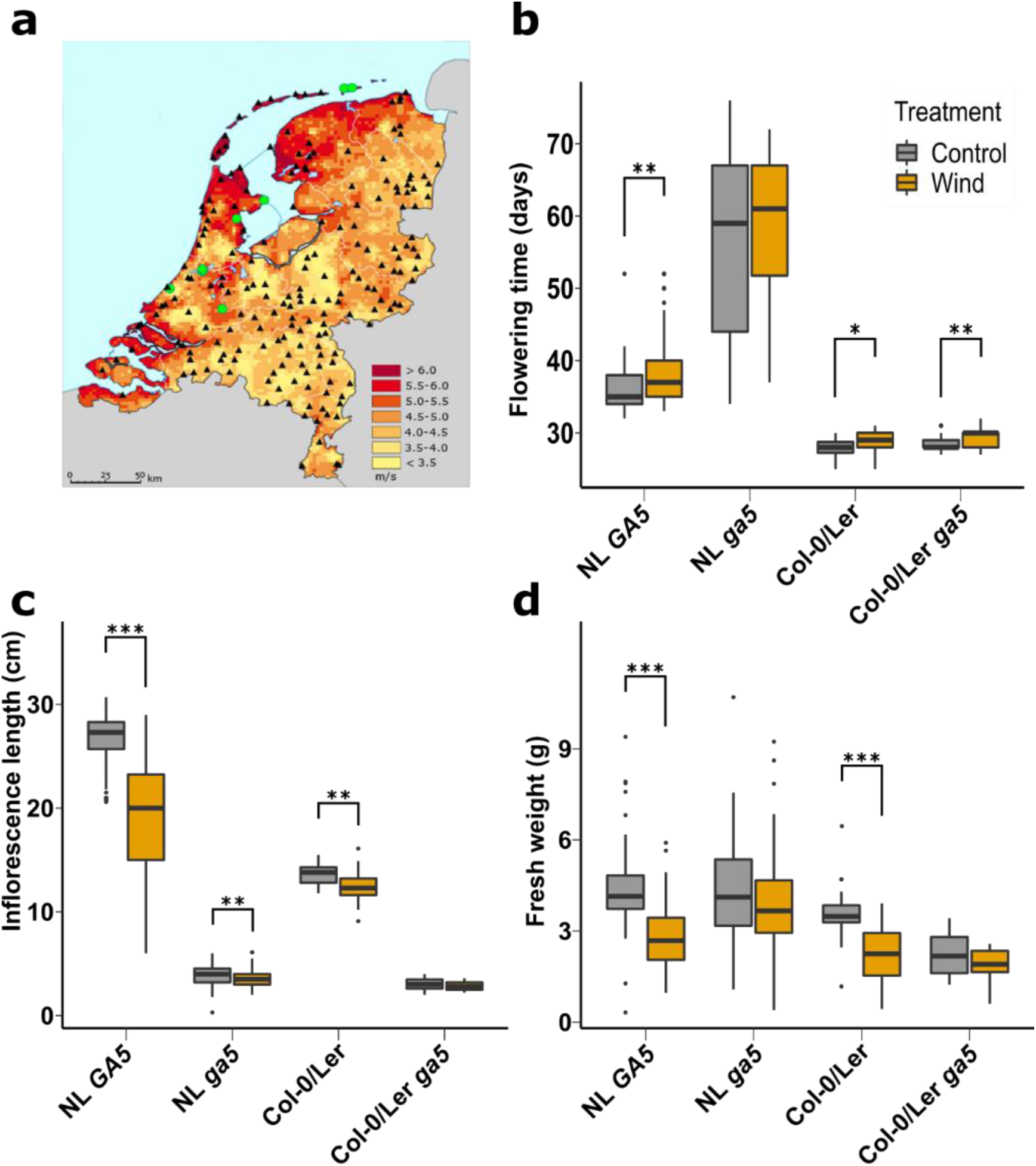
Phenotypic response to windy conditions. (a) Average annual wind speed (between 1981-2010) in the Netherlands as measured at ground level. Green circles indicate semidwarf accessions that share the same splice site mutation, black triangles indicate all other DartMap accessions. Figure adapted from https://www.knmi.nl/klimaat-viewer/kaarten/wind/gemiddelde-windsnelheid/jaar/Periode_1981-2010. (b-d) Plant responses to absence or presence of windy conditions (grey and orange respectively) for (b) flowering time, (c) inflorescence length one week after the first flower opened, (d) fresh weight at flowering \**P <* 0.05; \*\**P <* 0.01; \*\*\**P <* 0.001, two-tailed Student’s *t*-test. NL – Dutch accessions; *GA5* – wild-type allele, *ga5* – mutant allele.

We grew six Dutch semidwarf accessions and five unrelated tall accessions (Table S4), in a climate-controlled growth chamber equipped to provide controlled wind speeds typically experienced in coastal areas of the Netherlands in March/April.

Moreover, we included Col-0, L*er* and the *ga5* loss-of-function mutants in Col-0 and L*er* backgrounds for a direct comparison of near isogenic lines differing only at the *GA5* locus. Despite our preselection for similar flowering time (Table S4), based on previous data collected on greenhouse-grown plants, the semidwarfs flowered significantly later in the controlled growth room conditions than the taller accessions (Figure 3b). This likely had little effect on our final results though, as all accessions were exposed to windy conditions for the same time and all plants started to flower before the end of the experiment. The tall accessions were strongly affected by the wind, as evidenced by a significant reduction in stem length (approximately 30% on average) and fresh weight (approximately 36% on average) (Figure 3c and 3d). In contrast, the semidwarfs also showed a significant reduction in stem length, but the average reduction was only around 10%. No significant decrease in fresh weight was observed (Figure 3d). Similar effects were observed for Col-0 and L*er*, where the wind treatment significantly decreased their stem length and fresh weight, whereas the respective *ga5* loss-of-function mutants remained unaffected (Figure 3c and d). Collectively, these results strongly suggest that the semidwarf phenotype caused by the *ga5* mutation provides a strong advantage with respect to plant biomass production, when growing in conditions of high wind speeds.

### Flowering time variants display a latitudinal cline

The transition from vegetative to reproductive stage is a key life history trait in *A. thaliana*, influenced by seasonal cues like day length and temperature (Andrés and Coupland, 2012). Previous GWAS of flowering time yielded few significant associations, mainly due to allelic heterogeneity (Atwell et al., 2010; Tabas-Madrid et al., 2018; Zhang and Jiménez-Gómez, 2020). We expect this should be less of a problem in a regional population with more closely related individuals. Therefore, we performed GWA analysis of flowering time, with and without vernalization, using the DartMap panel. We discovered a highly significant association between flowering time and a 16-bp indel near the end of the first exon of the *FRIGIDA* (*FRI*) gene (position 269960 on chromosome 4 of Col-0), irrespective of vernalization treatment (Figure 4a). This indel is known to cause a premature stop codon in the *FRI* coding sequence, leading to a non-functional *FRI* allele (Koornneef et al., 1994; Johanson et al., 2000; Andrés and Coupland, 2012). *FRI* is a key regulator of flowering time in *A. thaliana* and loss-of-function alleles are widespread in nature (Zhang and Jiménez-Gómez, 2020). As the variant we picked up by GWA analysis is unlikely to be the only loss-of-function mutation to occur in the Netherlands, we examined all polymorphisms in the *FRI* coding sequences (CDS) in the DartMap and found a total of 41 *FRI* alleles. When compared to the *FRI-H51* allele, that encodes a fully functional protein and is regarded as the ancestral *FRI* allele based on sequence comparisons with *A. lyrata* (Le Corre et al., 2002; Zhang and Jiménez-Gómez, 2020), another 11 loss-of-function *fri* alleles are identified across 88 accessions. The accessions with a loss-of-function allele are on average early flowering (Figure 4b).

**Figure 4:**
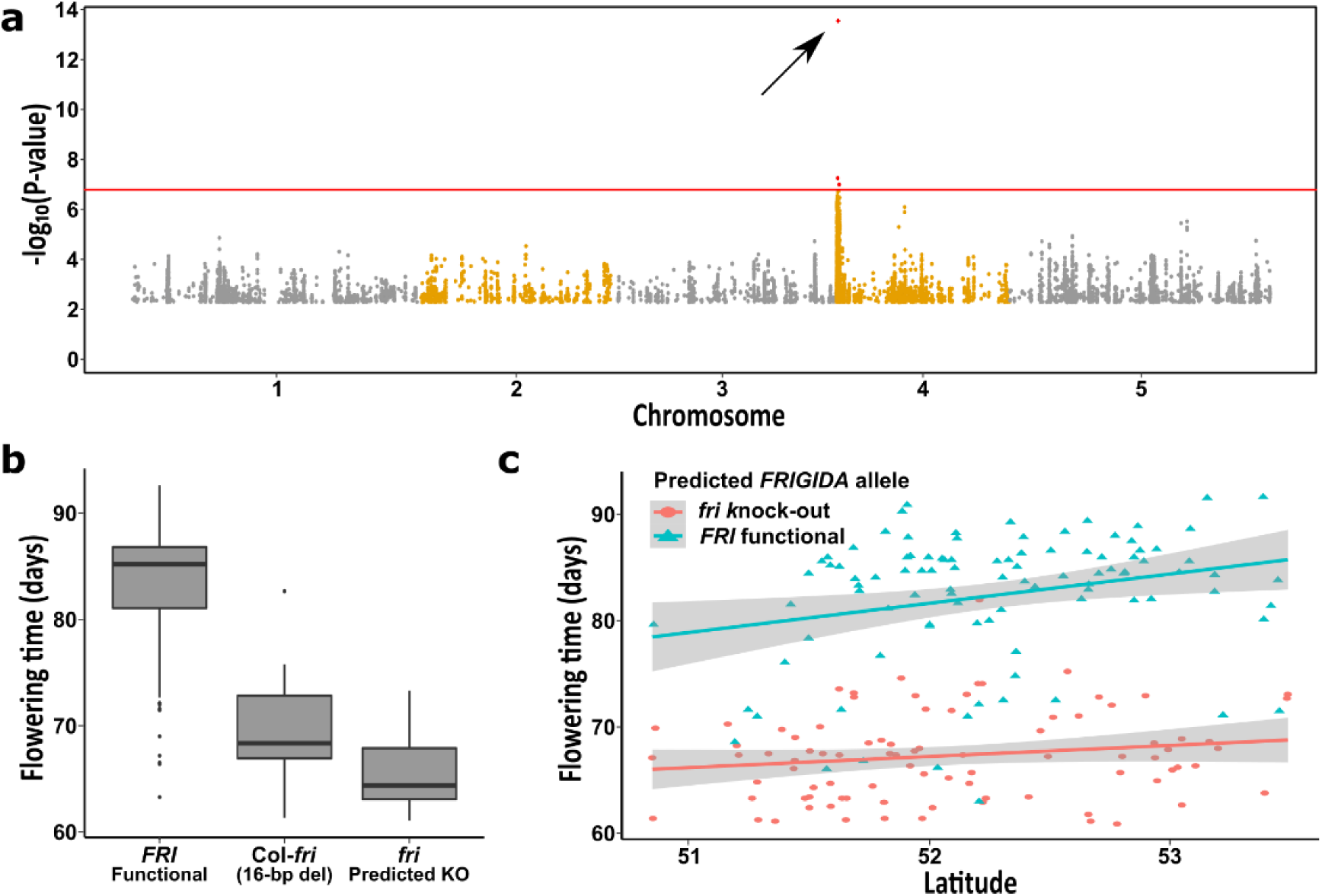
GWA analysis of flowering time in the DartMap panel upon vernalization. (a) Manhattan plot for days until flowering after a six-week vernalization treatment, identifying one highly associated polymorphism located in the *FRIGIDA* gene (arrow).

Although the Netherlands has a small latitudinal range, we examined if there is a latitudinal cline in the diversity panel regarding flowering time, consistent with previous findings in Northern European and Mediterranean accessions (Stinchcombe et al., 2004). While we did not observe such a cline when considering the entire population, when excluding the accessions carrying the *fri* loss-of-function alleles, we indeed found such a latitudinal cline (Figure 4c). This means other loci, next to *FRI*, are involved in controlling flowering time in the DartMap collection. These analyses collectively demonstrate the suitability of the DartMap panel for conducting GWA analyses on phenotypic variation relevant to local adaptation.

The red line corresponds to a permutation-based threshold level of significance at *P <* 0.05. (b) Flowering time distribution of three *FRIGIDA* (*FRI*) variants in the DartMap panel. Accessions with a functional *FRI* allele (*FRI F*unctional), accessions with the Col-0 variant (Col-*fri*) that corresponds to a 16-bp deletion causing a premature stop codon in the *FRIGIDA* coding sequence and accessions with a predicted functional knock-out allele of *FRI* (*fri* Predicted KO). (c) Average days until flowering for DartMap accessions relative to their latitudinal site of origin with linear fit for accessions with a predicted loss-of-function *fri* allele (red, R^2^ = 0.016, *P* = 0.128) or a functional *FRI* allele (blue, R^2^ = 0.082, *P* = 0.002).

### Local adaptation to the Dutch climate

The Dutch climate is characterized as a temperate maritime climate, which is relatively uniform across the Netherlands. However, there are mild and gradual clines in environmental variables, with slightly higher temperatures and less precipitation in inland areas compared to coastal areas. One advantage of having a densely and uniformly sampled population is the potential to detect whether local adaptation to these gradual environmental clines has influenced genetic diversity. To test this hypothesis, we conducted a genome-environment association (GEA) analysis on 19 quantitative climatic variables related to temperature and precipitation (Table S5 and Figure 5a as example). We identified a few QTLs for these variables, but the most notable QTL is located on Chr. 4 (Figure 5b). We named it *qDPT* (QTL for the Dutch Precipitation and Temperature climatic variables). This QTL is associated with seven climatic variables reflecting aspects of temperature and precipitation (Table S5).

**Figure 5:**
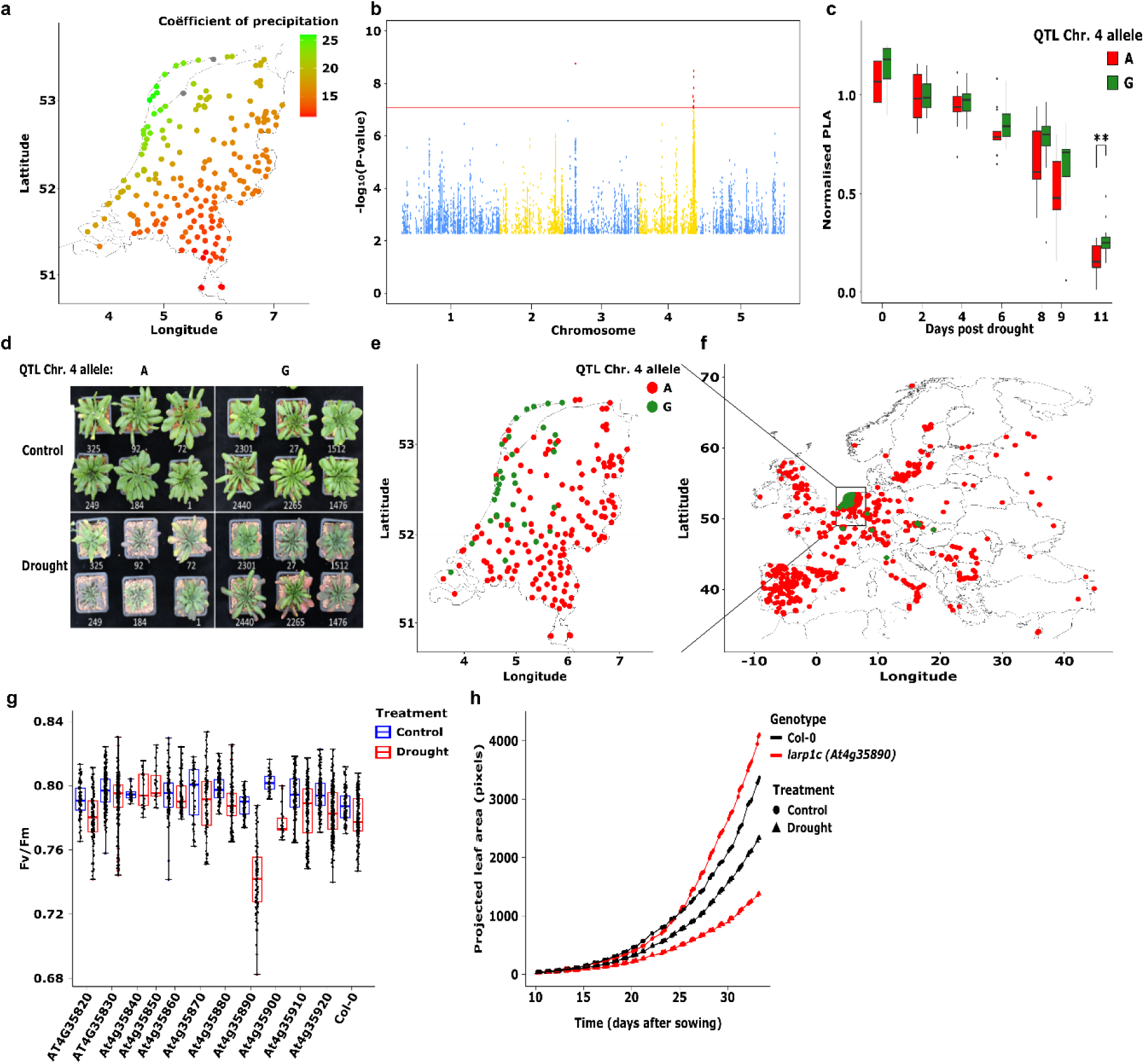
Genome-environment association (GEA) analysis on climatic variables related to precipitation and temperature. (a) Geographical distribution of precipitation seasonality (coefficient of variation) expressed in Seasonality Index (SI) units for the DartMap. SI values <0.19 are defined as ‘precipitation throughout the year’, whereas values between 0.20 and 0.39 are defined as ‘precipitation throughout the year, but with a definite wetter season’. (b) Manhattan plot for the GEA related to precipitation seasonality (coefficient of variation). (c) Response to the drought treatment for accessions differing at their *qDPT-B* allele. Boxplots represent the normalised projected leaf area (PLA) calculated per accession by dividing the PLA in the withholding watering treatment over the PLA in the control treatment for a set of 14 randomly chosen accessions per allele (depicted as an adenine (A) or guanine (G) nucleotide at SNP 4^17000635^). (d) Pictures of six representative accessions per *qDPT-B* allele in the control and drought treatment, taken 11 days after the start of the treatment. (e) Geographical distribution of the two alleles (‘A’ and ‘G’) across the Netherlands and (f) Europe. (g)) *Fv*/*Fm* as a proxy of plant stress of Col-0 and candidate gene T-DNA insertion lines. (f) The response of PLA of Col-0 and the *larp1c* mutant in the control and drought treatment.

Further examination of *qDPT* revealed that it appears to consist of two separate QTLs, approximately 230 kilobase pairs apart (Figure S6a). The SNPs that are found most frequently with the highest association in each QTL are SNP 4^16770900^ and SNP 4^17000635^ with minor allele frequencies (MAF) of 14.2% and 17.5%, respectively. To determine whether the -log(*P*) peaks represent distinct QTLs or a single one, we calculated pairwise *r*^2^ values between SNP 4^16770900^ and all other biallelic SNPs in the selected genomic region (16,700,000–17,200,000 bp). This revealed only weak evidence of linkage disequilibrium (LD) (pairwise *r*^2^ = 0.52) between the two SNPs most strongly associated with the trait (Figure S6b). Hence our conclusion these are two separate QTLs. For convenience, we will refer to them as ‘*qDPT-A*’ and ‘*qDPT-B*’, with their corresponding SNPs labelled as A/a for SNP 4^16770900^ and B/b for SNP 4^17000635^.

The GEA analyses provide insights into whether specific alleles are correlated with particular climatic variables. A strong association at a specific locus can indicate that allelic variation at that locus is important for local adaptation (Ferrero-Serrano and Assmann, 2019). However, such a strong correlation could also arise by accident, if the climatic variable happens to align with the population structure. We confirm that this is not a case of such a potential statistical artifact. First of all, kinship correction is used to perform the GEA analyses, and this should drastically reduce the chance of finding loci that are strongly associated due to the population structure. Secondly, we observe that the geographical distribution of the accessions contributing to the population structure appear to be mostly affected by the river systems (Figure 2b), which can be regarded as a South – North cline. The climatic variables that map at *qDPT* follow a mostly East – West cline. Moreover, the minor allele of *qDPT-A* is found in five out of six nuclear subclusters (the common allele is found in all subclusters), and for *qDPT-B* both alleles are found all subclusters. In conclusion, identification of these QTLs is not an artifact due to the general population structure and thus likely to represent a locus associated with plant adaptation to the environment.

It is not immediately apparent to which specific climatic variable(s) this locus contributes. The climate in the Netherlands is primarily influenced by the North Sea, which affects various environmental factors such as temperature, wind speed and precipitation patterns and variability. Therefore, it is not surprising that the same peak is consistently observed for several climatic variables, given the high correlation among the variables associated with *qDPT*. Consequently, unveiling the particular climatic variable or combination of variables to which the adaptation is associated with this specific locus may be challenging. However, we noted that the climatic variables associated to *qDPT* are all connected to seasonality, while annual means in precipitation and temperature are not associated. This suggests that the adaptation is related to conditions that prevail during certain parts of the year, such as one or two seasons, or to adaptations that enable the plant to effectively tolerate greater variability. To explore which climatic factor may be involved in this regard, we tested if allelic variation at *qDPT* resulted in a differential response to a gradual decrease in water availability, which is likely to be seasonal.

As *qDPT-A* and *qDPT-B* are in close proximity, we selected the best possible set of accessions distinguishing best the combinations of alleles for these QTLs aiming to disentangle the individual contributions of the QTLs. Given the available genotypes of accessions in the DartMap population, this results in 13 accessions with ‘AB’, 7 with ‘Ab’, 1 with ‘aB’ and 7 with ‘ab’ (Table S6). Initially, all accessions were grown under well-watered conditions. Subsequently, after two weeks, half of the replicates for each accession were no longer watered for eleven days. We refer to this as the drought treatment. We regularly measured the projected leaf area (PLA) during this period in the control and the drought treatment. Due to an uneven distribution of accessions across the genotypic combinations for these QTLs, we compared accessions with ‘Ab’ genotypes against those with ‘ab’ genotypes (7 accessions per group) to assess the effect of *qDPT-A*. For *qDPT-B*, all 28 accessions (14 accessions per *qDPT-B/b* allele) were included in the analysis. Our results indicated a significant effect of *qDPT-B* on the PLA response to the drought treatment (two-way ANOVA with *qDPT-B/b* allele and days after last watering as variables; *P* = 0.003) (Figure 5c and 5d). However, we did not observe a significant effect for *qDPT-A*. Subsequently, we examined the distribution of *qDPT-B* alleles across the Netherlands and the 1001G panel. We found that SNP 4^17000635^ (*qDPT-B*) has a much higher MAF in the Netherlands (MAF = 17.5%) (Figure 5e) compared to the 1001G panel (MAF = 3.1%) (Figure 5f).

We propose that the region encompassing the genomic locus with a LOD > 6, which is consistently observed across different mappings, is the most likely location for the gene(s), allelic variation of which causes the QTL. This region comprises 13 genes spanning from At4g35820 to At4g35940 (Table S7). Within this region, there are a few genetic variants that are in linkage disequilibrium (*r*^2^ > 0.4) with SNP 4^17000635^ and that potentially impact the encoded protein sequence of the gene they reside in.

Noteworthy are two non-synonymous variants in *OSCA4.1* (At4g35870), also known as *EARLY-RESPONSIVE TO DEHYDRATION STRESS 4* (*ERD4*). The first variant constitutes a change from methionine (neutral-polar) to isoleucine (hydrophobic) at the 10^th^ position, while the second variant changes an isoleucine to leucine (both hydrophobic) at the 641^st^ position. These substitutions constitute relatively minor sequence differences that are unlikely to affect the functionality of the protein encoded by the *ERD4* gene. We tested T-DNA insertion lines for 11 of the 13 candidate genes in response to control and drought treatments and measured *Fv*/*Fm* as a proxy of plant stress (Maxwell and ^J^ohnson, 2000^)^ (Figure 5g). For the remaining two genes, At4g35930 and At4g35940, no T-DNA insertion mutant lines were available. The mutant of *LARP1c* (At4g35890) has significantly lower *Fv*/*Fm* values in the drought treatment compared to Col-0, while it does not differ from Col-0 in the control treatment (Figure 5g). Furthermore, the mutant PLA is slightly higher, though not significantly different from Col-0 PLA under the control treatment, but it is significantly impaired under the drought treatment (Figure 5h). These results pinpoint *LARP1c* as the most prominent candidate gene to underlie the difference in response to drought.

### Natural allelic variation for *FSD3* affects iron deficiency tolerance

Although the Netherlands is relatively uniform in climate, the soil type composition differs substantially across the country, with clay, peat and sand forming the major types (Hartemink and Sonneveld, 2013). This could be an important driver for local adaptation in the Netherlands, as nutrient composition and especially nutrient availability (Veer, 2006), differs between soil types. Iron (Fe) in particular is known to vary in availability and shortage of Fe will severely limit plant growth (Hell and Stephan, 2003). We determined the light-use efficiency of photosystem II (PSII) electron transport (Φ_PSII_) of the DartMap diversity panel grown under control and Fe-deficient conditions, through chlorophyll fluorescence measurements. Low Φ_PSII_ can serve as an early indicator of Fe deficiency (Terry, 1980), as Fe is an essential component of iron-sulphur, and heme proteins that play pivotal roles in photosynthesis and respiration (Hänsch and Mendel, 2009).

The low Fe supply leads to only mild visible symptoms, such as reduced plant growth and mild leaf chlorosis, but causes extensive variation for Φ_PSII_, much more than in the control condition (Figure 6a). GWA mapping identified a single associated region, on chromosome 5, with the most significantly associated SNP located at position 7,856,132 (Figure 6b). The same locus, and the same SNP, is also found when the average Φ_PSII_ values in the Fe-deficient condition, and the ratio between Φ_PSII_ in Fe deficiency versus control, are used as traits for GWA mapping (Figure S7). To identify candidate genes for the causal allelic variant, we limited the associated region to 22 kb, based on the genome positions of SNPs that are in linkage disequilibrium (LD) (at r^2^ > 0.55) with the most significantly associated SNP. This region is predicted to contain 10 genes (Figure 6c).

**Figure 6:**
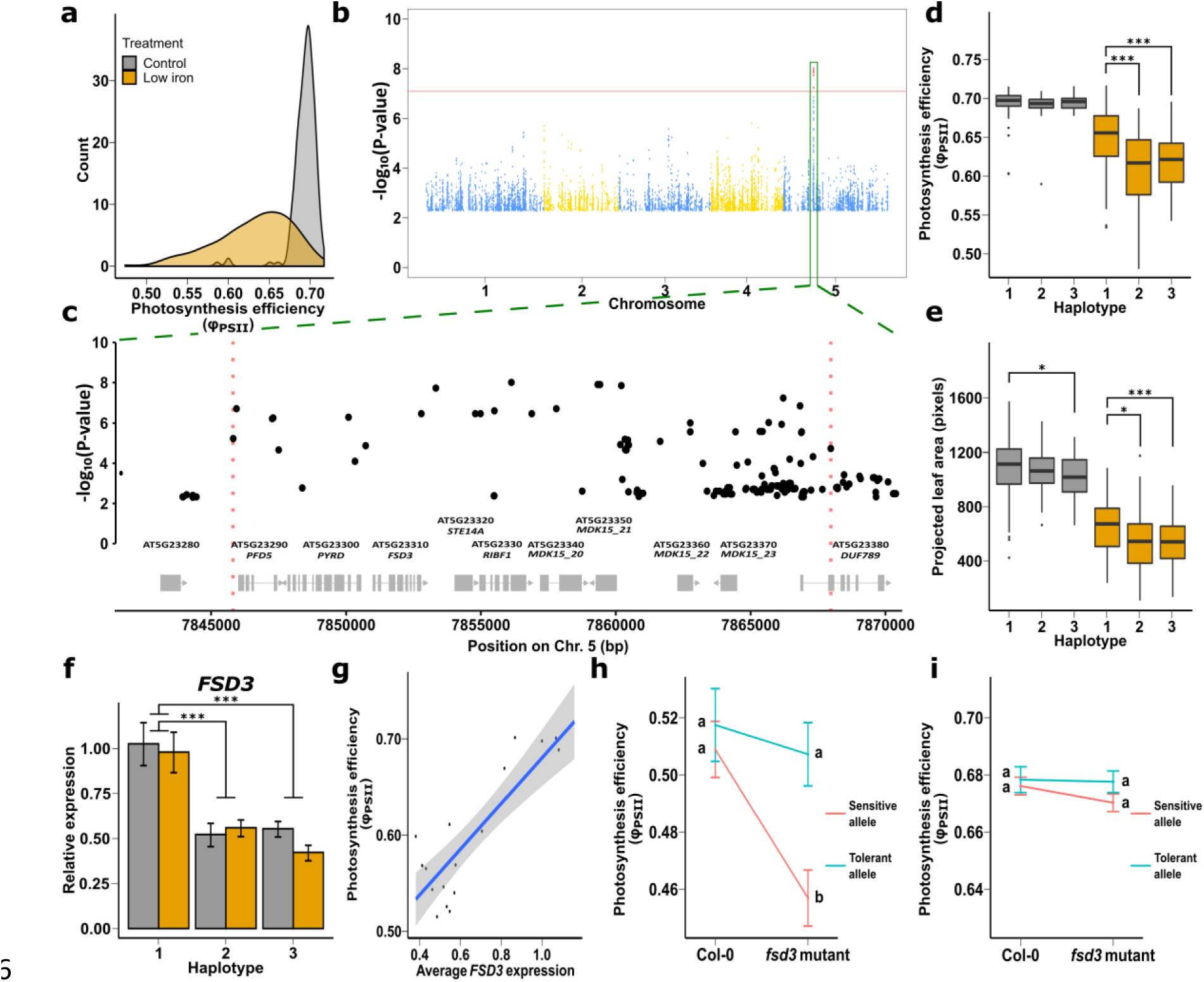
Genome Wide Association (GWA) analysis of the response of photosynthesis efficiency to iron deficiency. (a) Distribution of Φ_PSII_, the light-use efficiency of photosystem II (PSII) electron transport, when grown in control (21 µM Fe^2+^, grey) and low iron supply (1 µM Fe^2+^), orange). (b) Manhattan plot of the SNP variant associations [-log(*P*) > 2] when using the residual values of the linear regression of Φ_PSII_ in control conditions versus iron deficiency conditions as trait for GWA analysis. The red line indicates a permutation-based threshold level of significance of *P <* 0.05. (c) The genomic region surrounding the significantly associated region on chromosome 5 (green box in (b)). Red dotted lines demarcate the area in linkage disequilibrium with the most significant SNP, containing candidate genes depicted below (in grey). (d-e) Phenotypic distributions for Φ_PSII_ (d) and projected leaf area (e) per haplotype. (f) Relative gene expression levels per haplotype for *FSD3*. (g) Photosynthesis efficiency relative to the average relative *FSD3* expression level per accession with a linear fit (*FSD3*: R^2^ = 0.74, *P* = 2.94 × 10^-6^). (h-i) Φ_PSII_ of F_1_ progeny of crosses between Col-0 (tolerant *FSD3* allele) or the *fsd3* knockout mutant with accessions homozygous for the tolerant allele (N = 5) (blue) and similarly with accessions homozygous for the sensitive allele (N = 9) (red) in iron deficiency (h) and control conditions (i). Error bars represent SE. \**P <* 0.05; \*\**P <* 0.01; \*\*\**P <* 0.001, two-tailed Student’s *t*-test.

We identified three common haplotypes for this genomic region in the DartMap panel. Accessions that share the haplotype with the Col-0 variant associated with SNP5^7856132^ are referred to as haplotype 1, whereas accessions containing the non-reference variant are further distinguished into two haplotypes that are referred to as haplotypes 2 and 3 (Table S8). Haplotype 1 associates with higher Φ_PSII_ in Fe-deficient conditions relative to the non-reference haplotypes, which do not significantly differ in Φ_PSII_ (Figure 6d). The higher Φ_PSII_ of accessions with haplotype 1 correlates with a significantly higher projected leaf area in Fe-deficient conditions than haplotypes 2 and 3 (Figure 6e), while these do not differ significantly for projected leaf area.

In total, 40 sequence variants are in LD with SNP5^7856132^ at a cut-off of *r*^2^ *>* 0.55, of which 27 are intergenic, 7 intronic and 6 exonic (Table S9). Only two of the exonic variants encode a non-synonymous amino acid substitution, but in both cases the substitutions are from a leucine to an isoleucine or vice versa. These are very similar amino acids, and neither of these substitutions, one in *RIBF1* (At5g23330), the other in *DUF789* (At5g23380), appear to substantially change the function of these proteins, as predicted by InterPro (Mitchell et al., 2019).

We thus hypothesized that non-protein-coding variants that affect either levels or timing of gene expression should be causing the observed QTL. We measured expression of each candidate gene by means of RT-qPCR in both control and Fe-deficient conditions in shoots of six accessions of each haplotype (Table S9). Two known marker genes for Fe deficiency in leaves, *FE SUPEROXIDE DISMUTASE 1* (*FSD1)* and *FERRIC REDUCTION OXIDASE 3* (*FRO3)* (Waters et al., 2012), indeed showed the expected lower (*FSD1*) or higher (*FRO3*) expression under the Fe deficiency treatment compared to control (Figure S8a). Two of the 10 candidate genes, *FE SUPEROXIDE DISMUTASE 3* (*FSD3)* and *PRENYLCYSTEINE ALPHA-CARBOXYL METHYLTRANSFERASE (STE14A)*, are about 2-fold higher expressed in accessions with haplotype 1 compared to accessions with either of the other two haplotypes (Figure 6f and Figure S8b). The Φ_PSII_ in Fe deficiency correlates well with expression of these genes (Figure 6g). None of the eight other candidate genes showed such response of expression between haplotypes and correlation with the Φ_PSII_ in Fe deficiency (Figure S8b).

Since the absolute expression levels of *STE14A* are very low, we focused on *FSD3* as the most likely candidate to underlie the QTL. Two distinct splice variants are described for this gene that influence functionality regarding chloroplast development (Lee et al., 2019). We measured the relative gene expression of each splice variant independently, but found highly similar splice variant ratios between the different haplotypes (Figure S8c), indicating that natural variation affects gene expression of *FSD3* only quantitatively and not through altering differential splicing. Analysis of the non-protein-coding sequences surrounding *FSD3* identified a variable (TTC)_n_/(GAA)_n_ microsatellite repeat directly downstream of the *FSD3* 3’ UTR, which lacks either two or four of such repeats in haplotypes 2 and 3 relative to haplotype 1. As there are no differences in either Φ_PSII_ or *FSD3* expression between haplotype-2 and-3 accessions we distinguished two alleles for the *FSD3* locus based on their functionality in Fe deficiency: an Fe deficiency tolerant allele, which has the Col-0 reference sequence, and a sensitive allele lacking two or four (TTC)_n_/(GAA)_n_ repeats. We subsequently confirmed the causal nature of the allelic variation at *FSD3* to explain the QTL using quantitative complementation (Weigel, 2012), in which we test if the difference in Φ_PSII_ between F_1_ progeny derived from crosses between accessions with a tolerant allele crossed with both Col-0 and the *fsd3* mutant, and F_1_ progeny derived from similar crosses with accessions carrying the sensitive allele. In Fe-deficient conditions, the F_1_ progeny derived from the *fsd3* mutant and accessions with a tolerant *FSD3* allele significantly outperform the F_1_ progeny derived from crosses with accessions with a sensitive *FSD3* allele (*P* = 0.046) (Figure 6h). This is not found under control conditions (Figure 6i).

## Discussion

Here we present the DartMap panel, a new biodiversity panel consisting of 192 Dutch *A. thaliana* accessions, and we demonstrate its potential for detecting genetic signs of local adaptation. The genetic variation observed in this panel largely falls within the reported diversity of the 1001G collection, which is expected for accessions collected in Northwestern Europe, outside the European relict regions (The 1001 Genomes Consortium, 2016). The genetic diversity is unexpectedly high considering the relatively small geographical area without remarkable environmental clines. It illustrates the high level of standing genetic variation that exists among local populations of this predominantly self-pollinating species (Bomblies et al., 2010; The 1001 Genomes Consortium, 2016). Despite, or perhaps due to, the absence of strong environmental clines, the DartMap panel is very suitable for GWAS. The genetic architecture underlying natural adaptation is strongly dependent on the local environment (Brachi et al., 2013; Fournier-Level et al., 2013). Consequently, adaptive loci may exhibit genetic or allelic heterogeneity in regions with contrasting climates, which hampers their detection through GWAS. A milder diversity in environmental conditions can help to alleviate this limitation, as exemplified by the successful identification of the well-known *FRI* gene, which controls flowering time variation. This gene was not previously detected by GWAS in other collections such as the Iberian collection (Tabas-Madrid et al., 2018) or the large 1001G panel (The 1001 Genomes Consortium, 2016), primarily due to allelic heterogeneity. As such, the DartMap should be regarded as a valuable complement to collections sampled from regions with contrasting climates, and not a replacement. With the availability of several regional diversity panels, it will be easier to identify such adaptive loci when taking into account that each region may favour its own adaptive variation.

Our finding that genes involved in disease resistance are overrepresented among those impacted by CNV is consistent with previous work: a similar phenomenon was found for defence-response genes in a global set of 1,301 natural *A. thaliana* accessions (Göktay et al., 2021) and this same category of genes was found to be enriched in regions highly rearranged between seven diverse *A. thaliana* genomes (Jiao and Schneeberger, 2020). These observations suggest that CNVs may act as a genomic response to biotic stress. The next step would be to investigate whether such structural variants correlate to variation in the prevalence and abundance of known pathogens of *A. thaliana* (Bartoli et al., 2018).

The prevalence of plants with short inflorescences and the predominance of a near-isogenic genotype at various dispersed locations, strongly indicate an adaptive advantage for *ga5* mutants in the coastal regions of the Netherlands. Allelic heterogeneity and strong selection of one successful, near-isogenic genotype represented multiple times in the DartMap panel was the reason we could not identify the *GA5* locus through GWAS, but only based on prior knowledge on possible causal loci determining plant semidwarfism (Barboza et al., 2013). Signatures of selection were previously identified in *GA5* among *A. thaliana* accessions collected globally (Barboza et al., 2013), but the environmental cue driving such selection remained elusive. Later (Luo et al., 2015) identified several *ga5* loss-of-function plants in two Alpine populations of *A. thaliana* and noted a decrease in plant height with increasing altitude although the adaptive advantage to high altitude environments specifically was not conclusively demonstrated. Our findings demonstrate that high-wind treatment significantly impacts the growth vigour of tall Dutch *A. thaliana* accessions, whereas it has no significant effect on the vigour of predominantly coastally located semidwarf plants. Arguably, we determined this for plants that are growing under optimal water and nutrient supply.

Under natural circumstances, with additional growth-limiting factors and probably periods with much stronger winds, this reduction in vigour in the tall accessions is likely to be more detrimental, and also likely to affect their fitness (Younginger et al., 2017) or fecundity, which we could not establish under our artificial windy conditions. The observed phenotypic responses in terms of biomass and stem length to windy conditions are similar between the natural Dutch accessions and the Col-0 and Ler reference strains, as well as their corresponding *ga5* loss-of-function lines. Considering that wind speed generally increases with altitude in mountainous areas (https://map.neweuropeanwindatlas.eu), the selective advantage of the Alpine semidwarfs is also likely to be due to tolerance to wind speed. The selective pressure seems high, given the two independent *ga5* alleles we found in the Netherlands, and others found elsewhere in the world (Barboza et al., 2013; Luo et al., 2015). It is remarkable that the near-isogenic genotype collected at several Dutch sites spread over such large distances, without any obvious means of transport. This is in contrast to a previous case of whole-genome hitchhiking, due to selection on herbicide tolerance, where the spread was clearly linked to transportation along railway tracks (Flood et al., 2016a). However, given the high population density of the Netherlands, we assume that human-mediated movement of soil and seeds would also be sufficient to achieve dispersal of the near-isogenenic *ga5* semidwarfs.

The presence of a North-South cline in flowering time and the association between windy conditions and inflorescence height highlight the importance of relatively mild environmental clines for local adaptation. These patterns can be detected in a uniformly and densely sampled population like the DartMap. This is further exemplified by the strong association on chromosome 4 related to the mild cline in seasonality, where the alternative allele of *qDPT-B* is only found at a high local abundance in the West of the Netherlands. However, it should be noted that such associations do not necessarily provide evidence of causality. Demographic processes or genetic drift can produce similar patterns to those created by selection (Rellstab et al., 2015). To further investigate whether this association represents a case of local adaptation, common garden experiments or reciprocal transplanting experiments can be conducted (Hancock et al., 2011; Rellstab et al., 2015; de Villemereuil et al., 2016; Terés et al., 2019). Nonetheless, the high local relative abundance observed in the West of the Netherlands compared to rest of the world, along with the differential sensitivity to drought, suggests the presence of an adaptive locus. However, it seems unlikely that this adaptation is solely attributed to the drought response measured in our experiments, as similar climate characteristics in other regions would also be expected to exhibit a higher relative abundance. Therefore, it is more plausible that a combination of different climatic factors and/or other unconsidered factors such as urbanization contribute to this adaptation.

Identifying the causal gene in genotype-environment association analysis can be challenging because allelic variation may affect tolerance to an experimentally measured trait that may not necessarily reflect the trait under natural selection. However, in our study we found that the mutant of *La-RELATED PROTEIN 1c* (*LARP1c*) is more sensitive to drought compared to other tested mutants and Col-0. *LARP1c* encodes a protein with a conserved La-motif domain, which is associated with RNA recognition and is widely found among eukaryotes (Bayfield et al., 2010). Moreover, the sequence variant with the highest statistical association in *qDPT-B* (SNP 4^17000635^) is located in the 3’ UTR of *LARP1c*. Although neither of these observations provide conclusive evidence on whether this gene is underlying the *qDPT-B*, we consider *LARP1c* as the most plausible candidate gene. In *A. thaliana, LARP1c* is known to be involved in seed germination and the regulation of leaf senescence, which can be induced by various environmental stresses (Zhang et al., 2012; Yan et al., 2023). Additionally, each of the three LARP1 (1a, 1b, 1c) proteins in *A. thaliana* localise at the processing bodies (Zhang et al., 2012; Yan et al., 2023) and are proposed to play a role in RNA processes in response to environmental perturbations. For instance, *LARP1a* is required for thermotolerance of *A. thaliana* to long exposure of moderately high temperatures, where it is involved in regulating the RNA decay machinery together with the cytoplasmic exoribonuclease XRN4 (Merret et al., 2013). However, the specific functions of *LARP1c* in plants are still relatively understudied, and there may be other roles yet to be explored (Yan et al., 2023). *LARP1c* may thus also be involved in the response to drought and possibly also other environmental factors.

In addition to the mild and gradual environmental clines, we also examined the panel for sensitivity to low iron availability, as we expected to find variation for this trait given the distinct differences in soil type across the Netherlands. This identified non-coding variation at *FSD3* to cause a major QTL affecting sensitivity to low iron supply. *FSD3* is long known to encode an essential chloroplast-localized iron superoxide dismutase. Its function is to protect developing chloroplasts against excessive damage by superoxide (Myouga et al., 2008). However, natural genetic variation in *FSD3* has not been previously observed. The QTL affecting sensitivity to low iron supply is influenced by subtle allelic variation in *FSD3*, specifically related to the number of repeats in a microsatellite region directly downstream of the transcribed region. This variation results in a two-fold difference in gene expression between contrasting alleles, which strongly correlates with photosynthesis efficiency under iron-deficient conditions. When plants are supplied with sufficient iron, there is no phenotypic difference between contrasting haplotypes. FSD3 is one of two chloroplast-localized iron superoxide dismutases, the other being FSD2 (Gallie and Chen, 2019). While both can form heterocomplexes and *FSD3* overexpression can partly complement the *fsd2* mutant phenotype (Gallie and Chen, 2019), they are not genetically redundant. The *fsd2* loss-of-function mutant displays a compact rosette phenotype, with pale green leaves, and the *fsd3* loss-of-function mutant is barely viable, with a very small and pale rosette (Myouga et al., 2008). Since iron is an essential co-factor for iron superoxide dismutases, a decrease in iron supply is likely to affect their function. Under the mild iron deficiency conditions we tested, this decrease in function was apparently just enough to affect photosynthesis efficiency significantly more in the genotypes with lower *FSD3* expression.

We found no reason to suggest strong selection for either of the two alleles. Since the FSD3 function is essential for proper chloroplast development and tolerance to reactive oxygen species generated by photosynthesis, lower *FSD3* expression is likely disadvantageous. The impact of lower *FSD3* expression may even extend beyond chloroplast development and photosynthesis efficiency. For instance, hydrogen peroxide, which is produced when superoxides are converted by superoxide dismutases, can function as a signaling molecule in the systemic response to iron deficiency (Le et al., 2016). Therefore, plants with lower *FSD3* expression may have a delayed response in detecting iron deficiency. Nevertheless, a large ionome survey of 1135 global *A. thaliana* accessions did not reveal any clear indications of local adaptation to iron deficiency (Campos et al., 2021). This suggests that establishing a direct link between allelic variation and local adaptation to iron deficiency may be challenging. Factors such seasonal variation in iron availability depending on precipitation or limited distribution and heterogeneity of the allelic variation could contribute to the difficulty in establishing such a link.

In conclusion, our work demonstrates that regions with relatively mild environmental clines can still harbor a wide range of adaptive genetic variants. This highlights the value of establishing regional collections as excellent tools for studying local adaptation of *A. thaliana*. Such a collection does not need to be limited to regions with strong environmental clines as were put forward in previous studies (Brachi et al., 2013; Frachon et al., 2018). Even a diversity panel from a relatively small geographic region with mild environmental clines, such as the Netherlands, provides enough variation to pick up small differences in flowering time and sensitivity to iron deficiency. Given the decreasing costs of whole genome sequencing, it becomes increasingly easier to characterize the genetic variation in such populations in great detail. Our study may serve as a blueprint of this approach, supporting the view that collections of wild plant species can be powerful systems to determine the genomic basis of natural adaptation (Weigel and Nordborg, 2015; Sollars et al., 2017; Monnahan et al., 2019).

## Methods

### Plant material

A set of 192 lines was selected from a large collection of around 2,000 natural accessions sampled throughout the Netherlands, with the aim to obtain a uniform geographical spread of accessions and generate a local diversity panel, the <u>D</u>utch *<u>Ar</u>abidopsis <u>t</u>haliana* Map (DartMap) panel, suitable for GWAS (Table S2). While accessions were selected from a larger set of about 2,000 collected individuals, the selected panel has a relatively lower density of accessions in the north of the Netherlands and in the very south, due to limited availability or collector capacity. Seeds collected from individual plants in the field were used to propagate each accession as a line for two generations through single-seed-descent upon self-fertilization in a greenhouse, before DNA isolation.

### DNA isolation

Genomic DNA was extracted from one or more inflorescences (open flowers and above) from a single plant per accession. The material was frozen in liquid nitrogen, ground to a fine dust and incubated in 300µL 2x CTAB buffer (2% CTAB, 1.4m NaCl, 100mm Tris, 20mm EDTA, pH 8) for 30 minutes at 65^◦^C. An equal volume of chloroform was added, mixed, centrifuged (3250 rpm for 15 min) and the supernatant was collected. The DNA was precipitated by adding an equal volume of ice-cold isopropanol, incubated overnight at -20 ^◦^C and centrifuged (3250 rpm for 15 min). The precipitate was washed twice with 70% ethanol and air dried. DNA was dissolved in milliQ water and treated with RNase (Promega) for 30 minutes at 37 ^◦^C.

### Genomic DNA sequence data

Genomic DNA of the DartMap panel was sequenced at GenomeScan B.V., Leiden. Libraries were prepared using the NEBNext Ultra DNA Library Prep kit for Illumina, according to described procedures. 500-700 bp insert size fragments were sequenced using a Illumina Hiseq 4000 device, aiming for at least 30x coverage of paired-end reads (2×151 bp) of each accession, and used to generate the alignment files necessary for calling genomic variants (see Note S1 for details). We additionally generated alignments for the DartMap panel combined with 11 Dutch accessions included in the 1,001 Genomes (1001G) Project (The 1001 Genomes Consortium, 2016) (Sequence Read Archive ID: SRP056687). This combined set of samples is referred to as the “DartMap + 1001G panel”, to clearly distinguish it from the DartMap panel containing accessions collected in this study only.

### Single nucleotide polymorphism (SNP) and indel calling

Genetic variants were called using a workflow based on the Genome Analysis Toolkit (GATK) Best Practices (Van der Auwera et al., 2013). Base quality scores of aligned reads were recalibrated using GATK (version 4.0.2.1) BaseRecalibrator with default parameters, using a set of variants called in a world-wide panel of 1135 *A. thaliana* accessions as known sites (The 1001 Genomes Consortium, 2016) (obtained from ftp.ensemblgenomes.org/pub/plants/release-37/vcf/arabidopsis_thaliana). SNPs and short insertions/deletions (indels) were called in each sample using GATK HaplotypeCaller, allowing a maximum of three alternate alleles at each site. Samples were jointly genotyped using GATK GenomicsDBImport and GATK GenotypeGVCFs with default parameters. This last step generated three different VCF files containing calls for the nuclear genome, the mitochondrial genome, and the chloroplast genome respectively. We filtered these sets to remove putative false positive calls (see Note S1 for details).

### Copy number variation (CNV) calling

We called CNVs in the DartMap panel, defined here as deletions, duplications, and insertions of at least 50 bp, by applying the Hecaton workflow (Wijfjes et al., 2019) to the alignment files of each accession. Accessions of the 1001G were excluded from this analysis, as they were sequenced using a different sequencing platform, which can strongly affect CNV calling: a linear model of genetic distance between accessions based on CNVs as a function of differences in coverage, sequencing technology, insert size, and read length (see Note S1 for details) explained 30% of the variation in the number of CNVs found in each sample (*R*^2^ = 0.3). CNVs called in each accession were genotyped and merged to generate a single VCF file as output (see Note S1 for details).

### Characterizing the impact of deletions

We detected deletions overlapping protein-coding genes (by at least 1 bp) using bedtools intersect. Genes were obtained from the TAIR10 genomic annotation of Ensembl Plants (release 40) and from the annotation of the improved *A. thaliana* mitochondrial assembly (BK010421.1). Gene function was considered disrupted if the associated gene overlapped with a deletion and at least 76 bp (more than half the length of a read) of the coding sequence lacked read coverage. Deletions were defined as having a moderate frequency if they were present in at least 48 samples (25%) of the DartMap panel. We defined disease resistance genes by first collecting all Uniprot entries linked to all *A. thaliana* reference genes using the Retrieve/ID mapping tool (https://www.uniprot.org/uploadlists/), excluding a small number of genes (53) for which no corresponding entries could be found. Genes were labeled as involved in disease resistance if the name of their protein product contained the terms “disease resistance” or “Disease resistance”. Enrichment tests were performed using Fisher’s exact test as implemented in the Python SciPy library (Virtanen et al., 2020) (version 1.3.1).

### Population structure analysis

We clustered samples based on bi-allelic sites missing calls in fewer than 10% of the samples and having a non-reference allele in at least 10 of them, to ensure that accessions are grouped on variation at the population level, rather than rare variants found in few individuals. For clusters based on mitochondrial and chloroplast variants, this criterium was relaxed to a minimum of two samples, as a relatively large number of sites would have been discarded otherwise. To generate a distance metric, we represented samples by binary vectors with 1s for homozygous variants and 0 for all other cases and computed the Hamming distance between them, i.e. the number of variant sites differing between samples, using the dist.gene function of the R package ape (Paradis and Schliep, 2019) (v5.3). Samples were clustered with the hclust function in R (R Core Team, 2013), using complete linkage as the agglomeration method, similar to analyses on the Swedish and Iberian collection (Long et al., 2013; Tabas-Madrid et al., 2018) (Figure S9). We cut trees into different numbers of groups using the R function cutree, to explore the geographical distribution of clusters formed at various hierarchical levels. Geographical distance between accessions was computed using the distance function of the geopy library (https://github.com/geopy/geopy) (v2.0.0) with default parameters and their geographical locations were plotted using the basemap library (https://matplotlib.org/basemap/) (v1.2.2).

### Principal component analysis

Principal component analysis of chloroplast variants called in the Dutch and global collection was performed using the SeqArray (Zheng et al., 2017) (v1.28.0) and SNPrelate (Zheng et al., 2012) (v1.22.0) libraries in R. To remove putative false positive calls, variants were excluded if they had a quality-by-depth (QD) score lower than 25, leaving a total of 4095 sites. Furthermore, we removed 10 accessions in which at least 50% of the variants were called as heterozygous, suggesting DNA contamination of the samples used for sequencing. Results were visualized using ggplot2 (Wickham, 2016) (v3.3.2). Three outlier accessions were excluded from the final figures to improve the visualization of Dutch accessions with respect to the global collection. Accessions were coloured based on their chloroplast haplogroup assigned in a previous study (Hsu et al., 2019). Haplogroups of accessions that were not included in that paper (i.e. the DartMap panel) were predicted with a 1-nearest neighbour classifier on the accessions that were, using the first two principal components as features.

### Phenotyping flowering time and stem length

For all experiments, seeds were sown on wet filter paper and stratified for 3 days at 4 C before they were germinated. Seedlings were grown hydroponically on rockwool blocks (Grodan Rockwool Group, 40 x 40 mm in size) pre-soaked in a nutrient solution designed for *A. thaliana* (1.7mm NH ^+^, 4.5mm K^+^, 0.4mm Na^+^, 2.3mm Ca^2+^, 1.5mm Mg^2+^, 4.4mm NO_3_^−^, 0.2mm Cl^−^, 3.5mm SO_4_^2–^, 0.6mm HCO_3_^−^, 1.12mm PO_4_^3−^, 0.23mm SiO ^2−^, 21 µM Fe^2+^ (chelated with 3% diethylene triaminopentaacetic acid), 3.4 µM Mn^2+^, 4.7 µM Zn^2+^, 14 µM BO_3_^3−^ and 6.9 µM Cu^2+^ at pH 7, EC 1.4 mS cm^-1^). Plants were grown in a greenhouse until 97 days after sowing. For each treatment, plants were equally divided over 20 plastic trays, with 76-78 plants per tray. Eight replicates per genotype per treatment were used, randomly assigned over four separate trays with two replicates per tray per genotype. Each tray consisted of a unique, randomly generated combination of genotypes. Trays were placed in a random order in the greenhouse, and were moved once every two weeks to a new position to minimize possible spatial effects in the greenhouse. A vernalisation treatment of six weeks was performed in a climate-controlled growth chamber set at 4 ^◦^C, with 12h/12h light/dark period, 70 µM m^-2^ s^-1^ irradiance and 70% humidity starting at day 20 after sowing. After vernalisation, plants were returned to the greenhouse. Flowering time and height up to the first silique were scored in the greenhouse, either with or without a vernalisation period of six weeks. Flowering time was determined as the day when the first flower had opened. Stem length up to the first silique was measured one week after the first flower had opened, as this trait remains stable thereafter (Barboza et al., 2013). To obtain the set of phenotypes for further analysis, we calculated the average phenotypic values per accession per treatment.

### Wind simulation treatment

In a separate experiment, plants of six semidwarf accessions and five tall accessions of the DartMap panel, with similar flowering time, as well as the accessions Col-0 and Ler, and their respective loss-of-function *ga5* mutants (Table S4) were grown for the first 14 days after sowing in a climate-controlled growth chamber, set at 20/18 ^◦^C (day/night) with 10 h light (from 08:00 - 18:00) and 70% humidity at an irradiance of 200 µM m^-2^ s^-1^ at ambient CO_2_ levels. The growth chamber contained two treatment basins. The treatment basins have the same design as described in a previous study (Flood et al., 2016b), with the distinction that the two treatment basins are physically separated and are on either side of the climate chamber. Each treatment basin was divided into five equally sized blocks to hold plants. Accessions were grown in a randomized complete block design with six replicates per block. Wind was simulated with two fans that were positioned at either end of one of the treatment basins. The fans were set to provide a wind speed that varied around 5.6 m/s for the majority of the treatment basin. The wind speed varied depending on the distance of the plant to the fan, with plants closest to the fan experiencing wind speeds up to 7 m/s and plants furthest away of the fan and at the edge of the treatment basin experiencing 3.5 m/s wind. The wind speed for the control treatment basin varied between 0.5 and 1.0 m/s. Wind speeds were measured with a TSI Velocalc (model: 8347-M-GB) air velocity monitor. Wind treatment was simulated by alternating patterns of six hours with wind by six hours without wind and was terminated when the first plants in the wind treatment basin had fully ripened siliques.

### Extraction of climatic variables

Climatic variables were obtained from google earth engine using R and RGEE (Aybar et al., 2020). All bands from Worldclim BIO data (Hijmans et al., 2005) selected from 250m-by-250m squares using the coordinates from the collection site for each accession as input.

### Drought treatment

To test which of the two nearby QTLs at *qDPT* affected drought tolerance, plants were grown in a greenhouse on ceramic-based granular soil. For the first two weeks, all plants were watered three times a week with the nutrient solution described previously. This watering regime was continued for the control treatment while for the drought treatment excess nutrient solution was removed and watering was ceased. Accessions (Table S6) were grown in a randomised complete block design with five replicates per treatment. Photographs were taken with a Nikon D3000 with the same settings, zoom and distance to the plants and were subsequently analysed to obtain the projected leaf area (PLA) with ImageJ (settings Hue 26-114, Saturation 58-218 and Brightness 121-255). The PLA was then normalised per accession by dividing the PLA in the drought treatment by the PLA in the control treatment to correct for innate differences in size of different accessions. For candidate gene validation T-DNA insertion lines were used (Table S7). Plants were grown in our phenotyping growth chamber, the Phenovator (Flood et al., 2016b), with the same conditions in the growth chamber as described for the wind simulation treatment. The drought treatment was induced in a similar way, by using ceramic-based granular soil. In contrast to the experiment in the greenhouse, the growth substrate was covered with black plastic cover to prevent algal growth that might interfere with high throughput phenotyping. Moreover, we watered plants in the drought treatment only once at the start, as there was less evaporation in this setup compared to the greenhouse. Starting at 10 days after sowing, we measured projected leaf area based on near infrared reflectance (NIR) imaging as described by (Flood et al., 2016b) four times per day, and F_v_/F_m_ using chlorophyll fluorescence once per day.

### Iron deficiency treatment

In a separate experiment, plants were grown as described for the wind simulation experiment, but in this case, seven rather than five blocks per treatment basin were used. Plants in the control treatment were watered with the nutrient solution described above, while those under iron deficiency treatment received an adjusted solution with only 1 µM Fe^2+^ instead of 21 µM Fe^2+^. Either three or four replicates were used per accession per treatment, with half of the accessions having four replicates in control condition and three replicates in the iron deficiency treatment and vice versa for the other half of the accessions. On day 15, plants were transferred to our phenotyping growth chamber, the Phenovator (Flood et al., 2016b), with the same conditions. We measured the steady state quantum yield of photosystem II electron transport (Φ_PSII_), as a proxy for photosynthesis efficiency, and the projected leaf area, based on near infrared reflectance (NIR) imaging as described by (Flood et al., 2016b), at five resp. seven time points per day. We discarded outlier samples with NIR values smaller than the mean minus two standard deviations per treatment. We corrected the raw phenotypic data (Φ_PSII_ and projected leaf area) for block effects by fitting the following linear mixed model in which random terms are underlined:

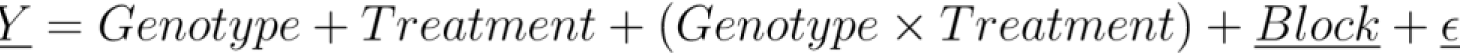

The model was analysed using the restricted maximum likelihood procedure with the lme4 package in R (Bates et al., 2015). For the GWAS, we calculated the average Φ_PSII_ per accession per treatment. To obtain a single phenotypic value representing a measure of tolerance per accession, we then plotted Φ_PSII_ for each accession under the iron deficient condition versus control condition and fitted a linear regression through the entire population. We calculated the residuals of each accession relative to this linear regression to be used as the phenotypic value for GWA analysis. Additionally, we used the average Φ_PSII_ values in the iron deficient condition and the calculated ratio between Φ_PSII_ in iron deficiency versus control as alternative traits for GWAS.

### Plant material for expression analysis

Plants that were used for gene expression analysis were grown in the greenhouse on rockwool blocks that were watered with either the control nutrient solution (21 µM Fe^2+^) or the adjusted low iron solution (1 µM Fe^2+^). For both treatments we used five plastic trays to contain the nutrient solution. Two replicate plants per accession were distributed over trays in a complete randomised block design. Whole shoots of two replicate plants per accession originating from the same tray were pooled and harvested as one sample for RNA isolation 21 days after germination, while five such samples per accession per treatment were harvested in total. The material was snapfrozen in liquid nitrogen immediately after harvesting and stored at -80 ^◦^C until RNA isolation. Accessions were selected for gene expression analysis (Table S8) based on their haplotype for the region with the highest association in the GWAS. Haplotypes were determined by looking at pairwise correlations between all variants with a minor allele frequency > 10% in a window of 20 kb surrounding the SNP with the highest association.

### Quantitative complementation

Quantitative complementation was used to test if the difference in Φ_PSII_ between the F_1_’s of Columbia-0 (Col-0) and the knockout mutant *fsd3* in Col-0 background crossed with accessions homozygous for the tolerant allele was significantly different from the response in Φ_PSII_ for F_1_’s of Col-0 and *fsd3* crossed with accessions homozygous for the sensitive allele. Five accessions with a tolerant, and nine accessions with a sensitive allele for *FSD3* were crossed to both Col-0 and *fsd3* (Table S8). Col-0 and *fsd3* were used as pollen donor in all crosses. The *fsd3* T-DNA insertion knockout mutant, SALK_103228, was ordered from the Arabidopsis Biological Resource Center and insertion of the T-DNA was validated by PCR as earlier described (Myouga et al., 2008). This T-DNA insertion line has no *FSD3* transcript (Myouga et al., 2008), and we could only grow it beyond the seedling stage on a 0.3% (w/v) gel-rite medium supplemented with 3% (w/v) sucrose.

Plants (N = 10 per cross per treatment) were grown as described in the section “Plant material for expression analysis”. Plants were phenotyped for Φ_PSII_ from 21 DAS onwards as described by (Baker and Oxborough, 2004) with the PlantScreenTM SC System (Photosystems Instruments). Phenotyping was done for twenty plants per run and lasted for three consecutive days overall. We phenotyped crosses with the tolerant allele and the sensitive allele equally spread out over the three days to prevent possible phenotypic differences caused by the delay between measurements. Moreover, in each measurement of twenty plants we included F_1_’s from both the cross between a given accession with Col-0 and with *fsd3* in a single measurement. We included one reference tray that was measured each day in the morning, at the start of the afternoon and at the end of the afternoon to test for possible differences in Φ_PSII_ during the day, but no significant differences were found. We corrected the phenotypic data for possible block effects by fitting a linear model, with random terms underlined:

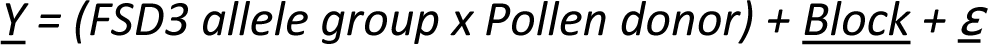

The model was analysed using the restricted maximum likelihood procedure with the lme4 package in R (Bates et al., 2015). We tested interaction between the *FSD3* allele group (natural tolerant or sensitive allele) and the pollen donor (Col-0 or the *fsd3* mutant) with a two-way ANOVA, for which a Kenward-Roger approximation for the degrees of freedom was used. Tukey’s post-hoc multiple comparison test (*α* = 0.05) was used to assess significance.

### RNA isolation and gene expression analysis

Total RNA was extracted with the Direct-zol RNA mini-prep kit (Zymo Research) according to the company’s instructions and then subjected to a DNAse (Promega) treatment at 37 ^◦^C for one hour. Total removal of DNA was validated by means of a no-reverse-transcriptase PCR reaction on a few mixtures of samples. The RNA quality was assessed for purity (A260/A280) with a spectrophotometer (Thermo scientific Nanodrop 2000) and for possible RNA degradation by means of a visual inspection of the RNA on a 1% (w/v) agarose gel. cDNA was then synthesized from 1 µg total RNA (measured by spectrophotometer) with the iScript^TM^ cDNA synthesis kit (Bio-RAD) according to the company’s instructions. Reverse transcriptase quantitative PCR (RT-qPCR) reactions were performed with the SensiFast SYBR Green Mastermix (Bioline) on a Bio-RAD cfx96 machine. RT-qPCR primer sets were designed in parts of the coding region that did not contain SNPs between the used set of accessions. For each RT-qPCR primer set used, we first validated its efficiency by means of a standard curve and only used primer sets with efficiencies ranging between 90% and 110%. Gene expression was normalized, using the delta cycling threshold (ΔCt) method (Livak and Schmittgen, 2001), to three reference genes, *SAND*, *YLS8*, and *TIP41-like* as earlier described (Han et al., 2013). Differential expression was assessed by two-sample t-tests on the ΔCt values. All primer sequences can be found in Table S10. We included *FSD1* and *FRO3* in the expression analysis as marker genes for iron deficiency in shoots (Waters et al., 2012).

### Genome-wide association (GWA) and genome-environment association (GEA) analyses

GWA and GEA mapping were performed in GEMMA (Zhou and Stephens, 2012) (version 0.98) with a univariate linear mixed model and the minor allele frequency cut-off set at 0.05. A kinship matrix was constructed in GEMMA to correct for population structure. CNVs were included in the analysis in a similar fashion as SNPs and indels by converting them into a reference variant, an alternative variant or heterozygous. To estimate a suitable genome-wide *p*-value threshold, we performed 1,000 permutations of the phenotype values. For each permutation we performed association analysis and extracted the highest association score. The distribution of these 1,000 highest association scores was then used to determine the empirical threshold value of *p <* 0.05.

## Data availability

Raw sequencing data of the DartMap panel can be found on the repository of the National Center for Biotechnology Information (NCBI) (BioProject ID: PRJNA727738).

## Authors’ contributions

SS, DDR and MGMA designed the project with RB and RYW. RB composed the panel. RYW performed computational analysis of sequencing data and overall patterns of genetic diversity. Wet-lab experiments were conducted by RB, FFMB, MM, JJV, NG and LMD. Subsequent data analysis of wet-lab experiments was performed by RB, MM, JJV, NG and LMD. TPJMT contributed especially to data collection and analysis of sequencing data of non-Dutch *A. thaliana* accessions. BLS extracted environmental data. RYW, RB, BLS, JJV, MM, NG, LMD, MK, FAvE, SS, DDR, and MGMA contributed to the experimental design, interpretation of results, and preparation of the manuscript. All authors read and approved the final manuscript.

## Acknowledgements

We thank Yanda Zhou, Jeroen van Buren, and Imme Bartels for their contributions to the data collection of phenotyping experiments; the Unifarm staff for taking excellent care of all plants grown during this work; and Korbinian Schneeberger for providing feedback on the manuscript. We are especially grateful to Erik Wijnker and Joost Keurentjes, who called upon the listeners of the Dutch radio program “Vroege Vogels” (https://www.bnnvara.nl/vroegevogels) to collect *A. thaliana* plants and seeds and send them to Wageningen University, and all those people who did. Without them, we would not have been able to get such a well-distributed set of Dutch *A. thaliana* accessions.

