## Supplemental Figures and Methods for "Allelic variants confer Arabidopsis adaptation to small regional environmental differences"

### Supplementary Figures

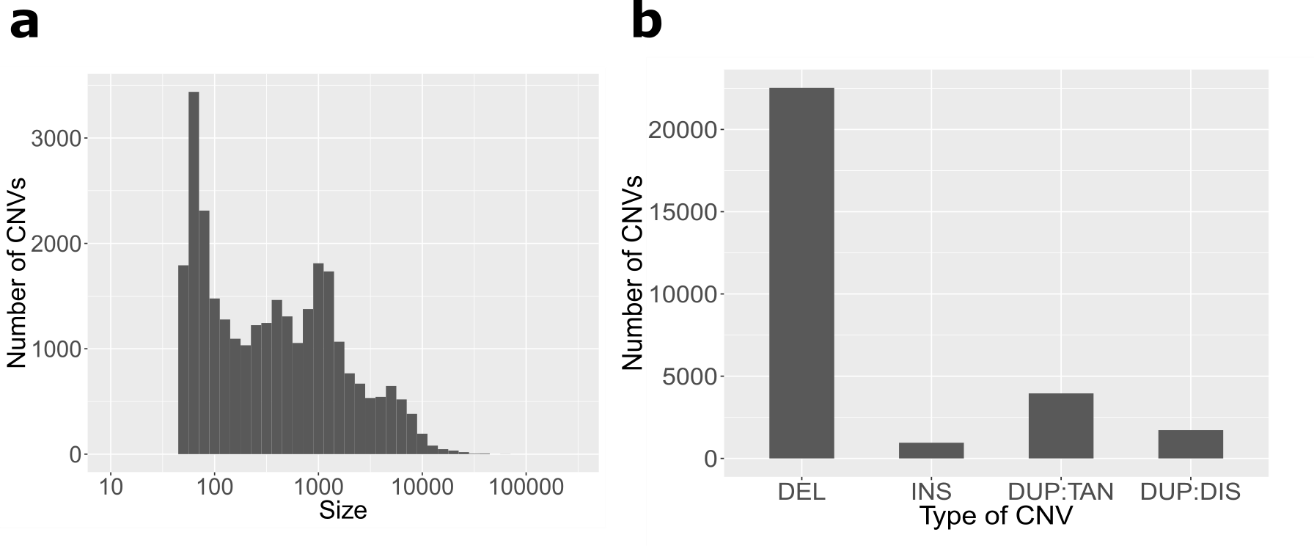
Figure S1: **CNVs in the DartMap panel.** (a) Size distribution of CNVs. CNVs below 50 bp are excluded, as these are considered indels. (b) Type distribution of CNVs: deletions (DEL), insertions (INS), tandem duplications (DUP:TAN), or dispersed duplications (DUP:DIS).

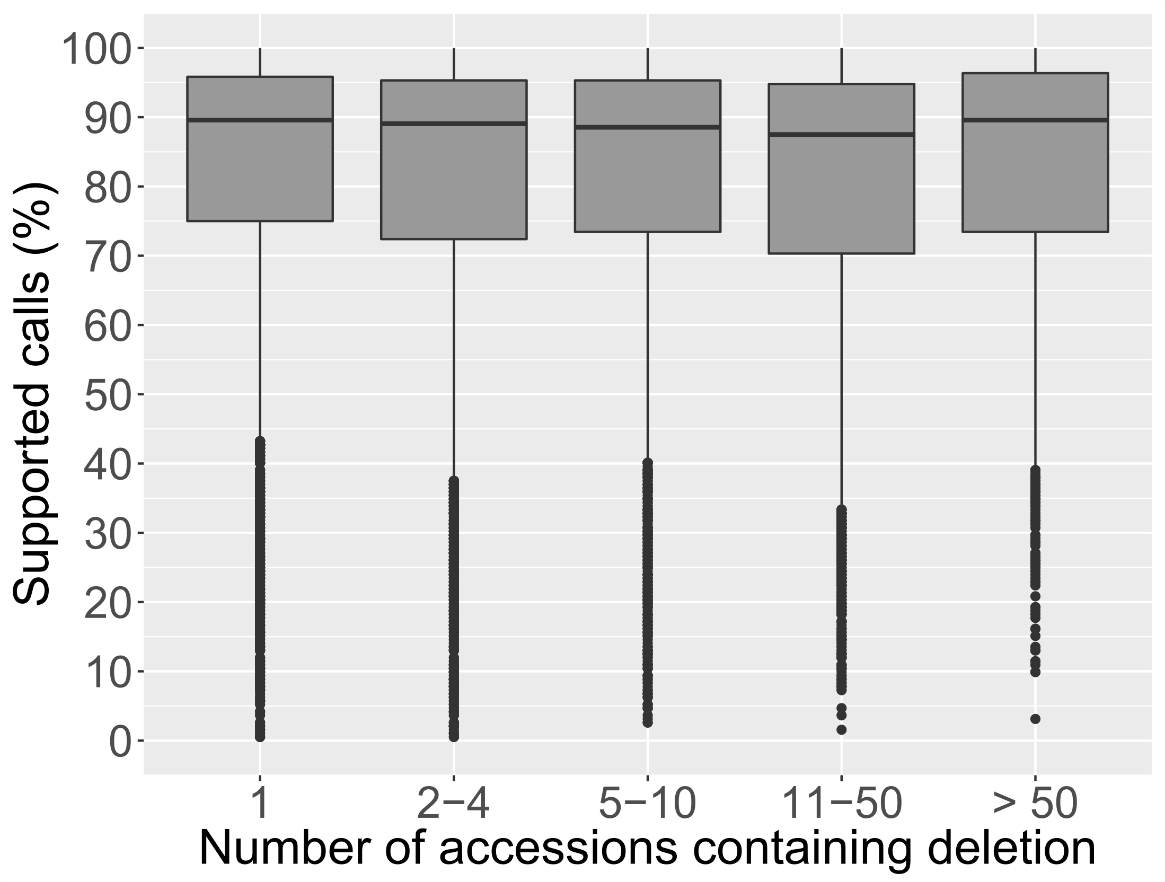

Figure S2: **Deletions in the DartMap panel were called with high precision.** The y-axis shows the percentage of genotype calls supported by a matching decrease (non-reference allele) or no change (reference allele) in read depth. Deletion variant sites are stratified along the x-axis by frequency of the non-reference (deletion) allele.

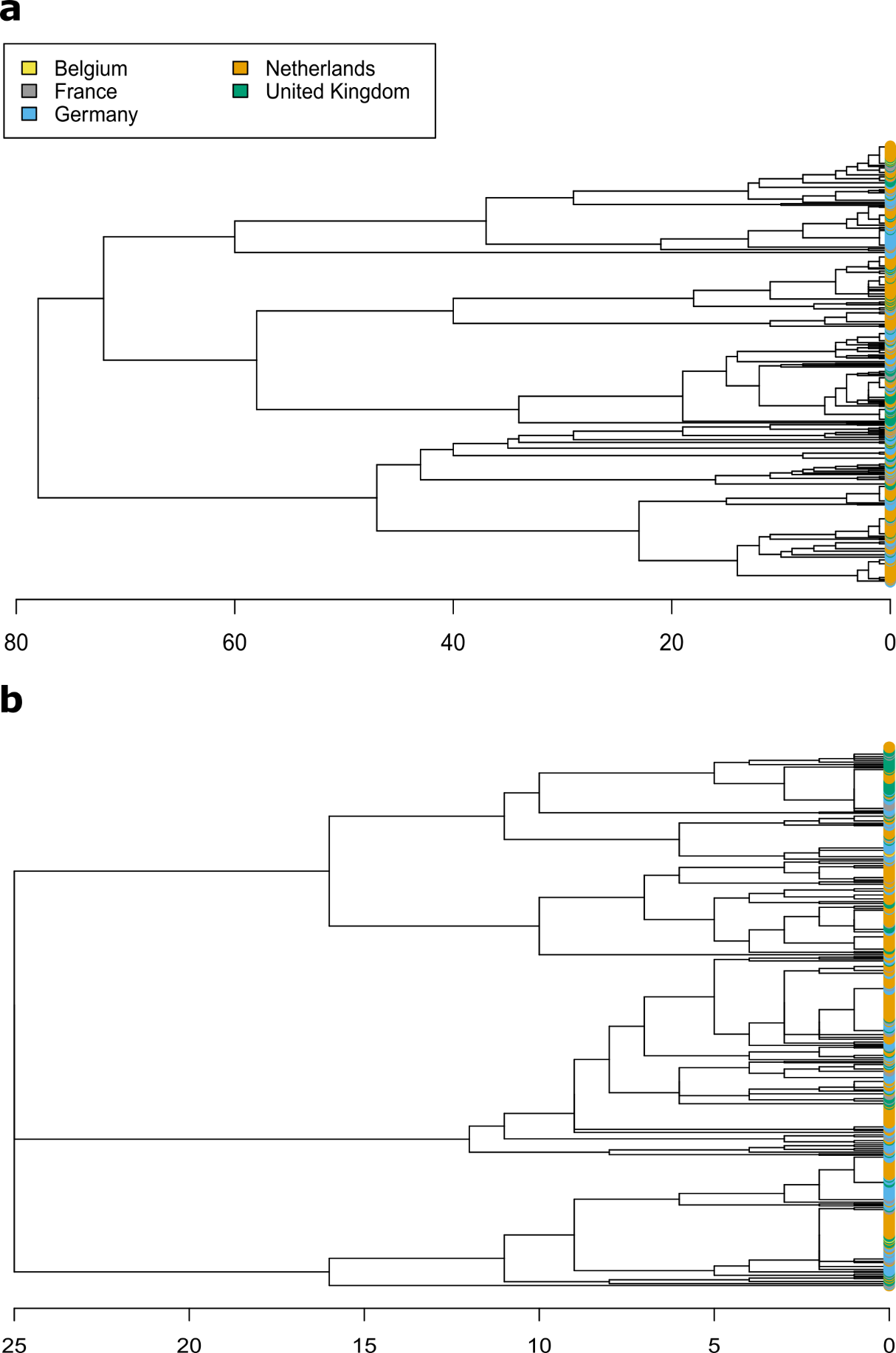

Figure S3: **Dendrograms based on organellar variation.** (a-b) Dendrograms constructed based on pairwise similarity of chloroplast (a) and mitochondrial (b) variants between Dutch accessions and those of nearby countries.

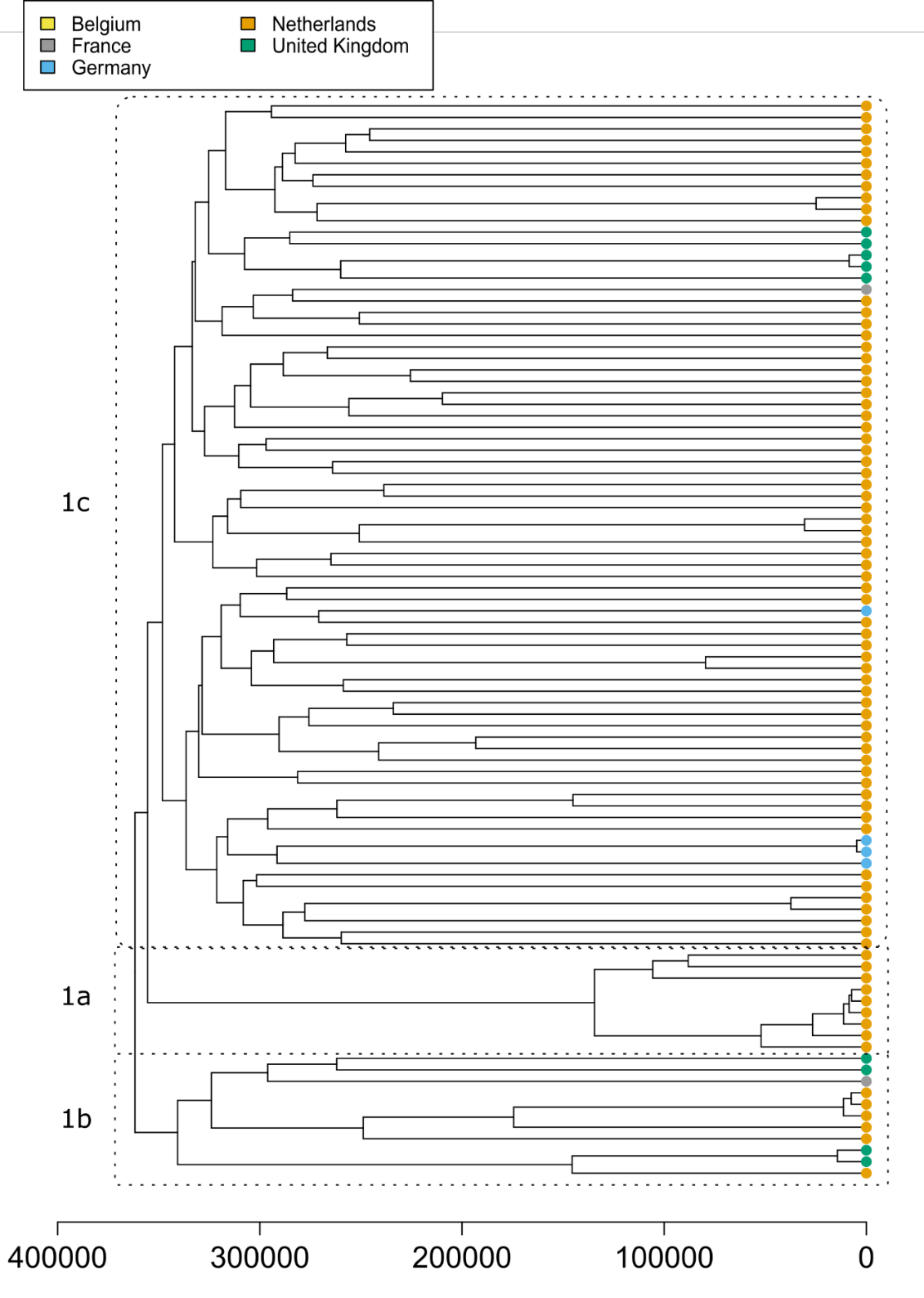

Figure S4: **Dendrogram of group 1 based on nuclear variation.** At its highest level, the dendrogram constructed based on pairwise similarity of nuclear variants between Dutch accessions and those of nearby countries can be split into two groups. Group 1 and its three subgroups (1a, 1b, and 1c) are depicted here.

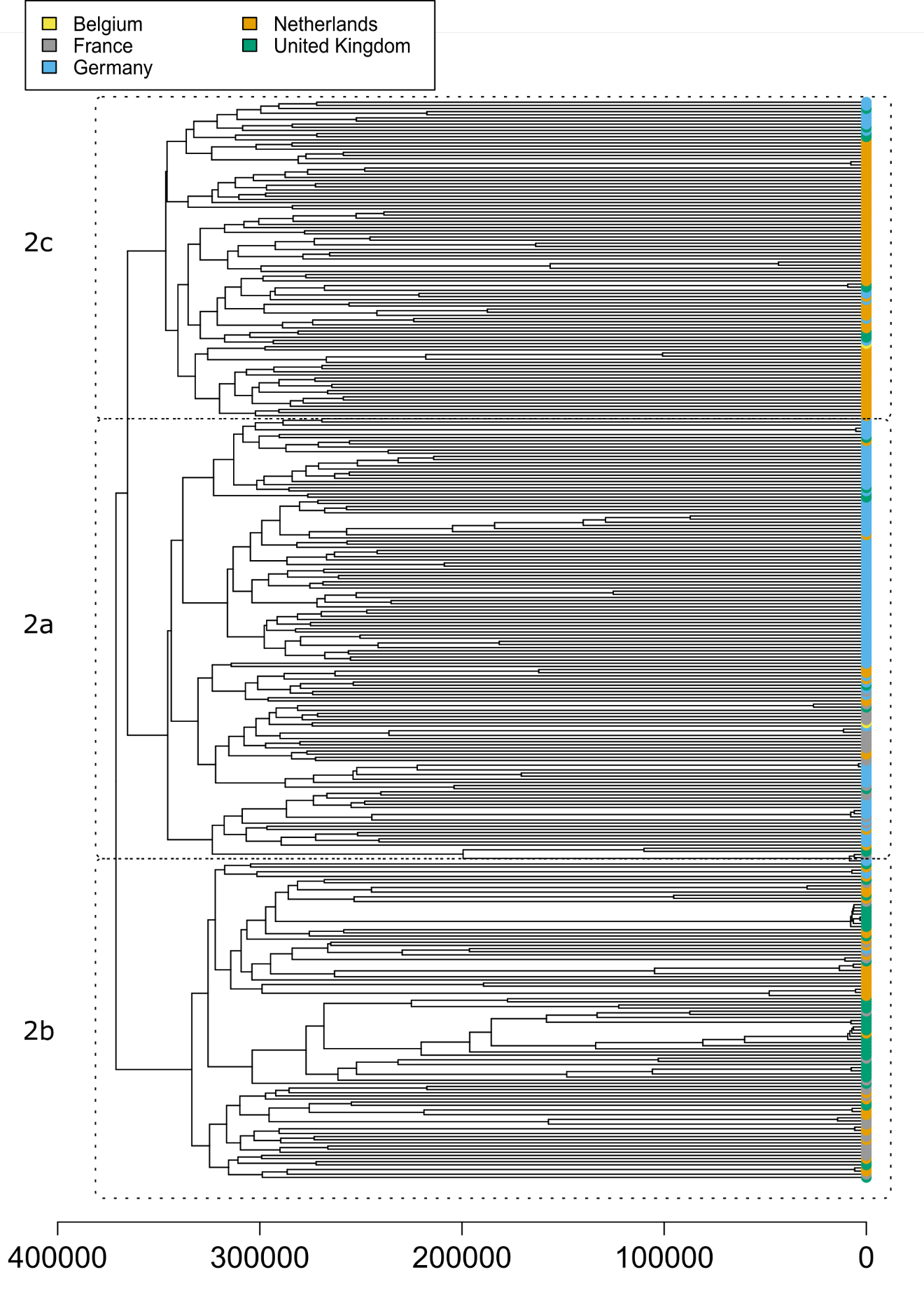

Figure S5: **Dendrogram of group 2 based on nuclear variation.** At its highest level, the dendrogram constructed based on pairwise similarity of nuclear variants between Dutch accessions and those of nearby countries can be split into two groups. Group 2 and its three subgroups (2a, 2b, and 2c) are depicted here.

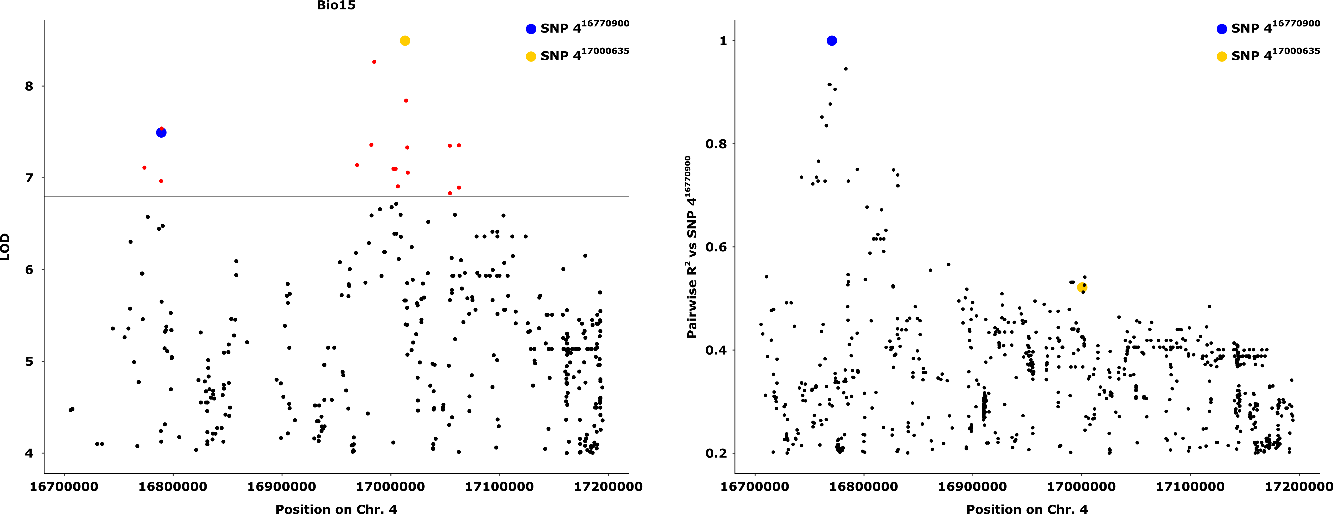

Figure S6: **Genomic region surrounding the chromosome 4 peak that was mapped for multiple climatic variables.** (a) A zoomed in section of the Chr. 4 peak of the mapping of precipitation seasonality (Figure 5b) shows that there appear to be two nearby QTLs. The two SNPs that were found most frequently with the highest LOD score among the different mappings are depicted with a blue and yellow dot. b) Linkage analysis (expressed as the pairwise R^2^) of SNPs in the Chr. 4 peak area relative to SNP 4^16770900^ (blue dot) reveals that the two QTLs are not in strong LD.

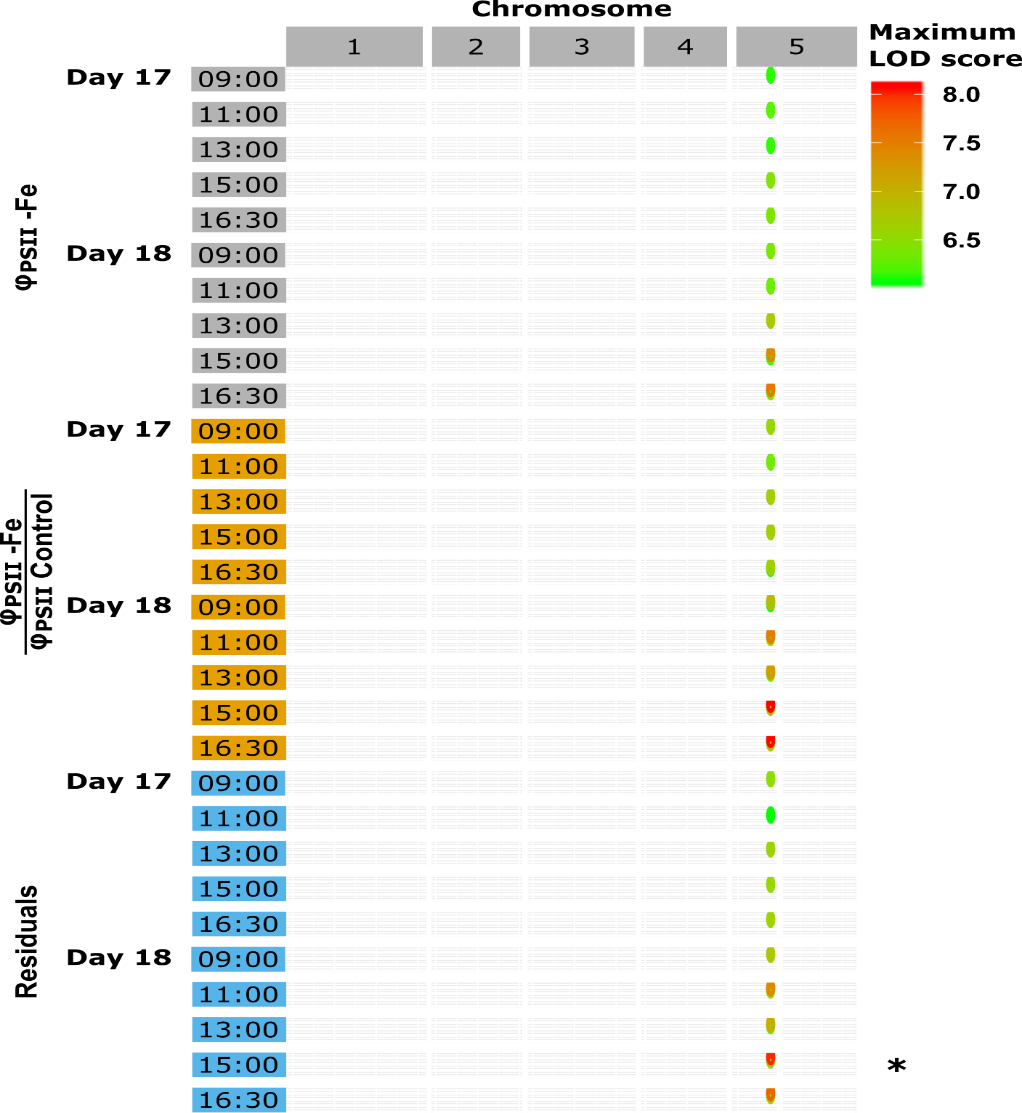

Figure S7: **Iron deficiency GWAS time series.** Summary data of the GWA results over 10 time points, measured during days 17 and 18 after sowing for three different phenotypes: [1] the average Φ_PSII_ per accession in iron deficient conditions (grey), [2] the ratio between Φ_PSII_ in iron deficient conditions and Φ_PSII_ in control conditions (orange), and [3] the residuals (blue). The dots on the graph represent the highest LOD scores in a 25 kb window size above a LOD threshold of 6. The * represents the data from the Manhattan plot in Figure 6b.

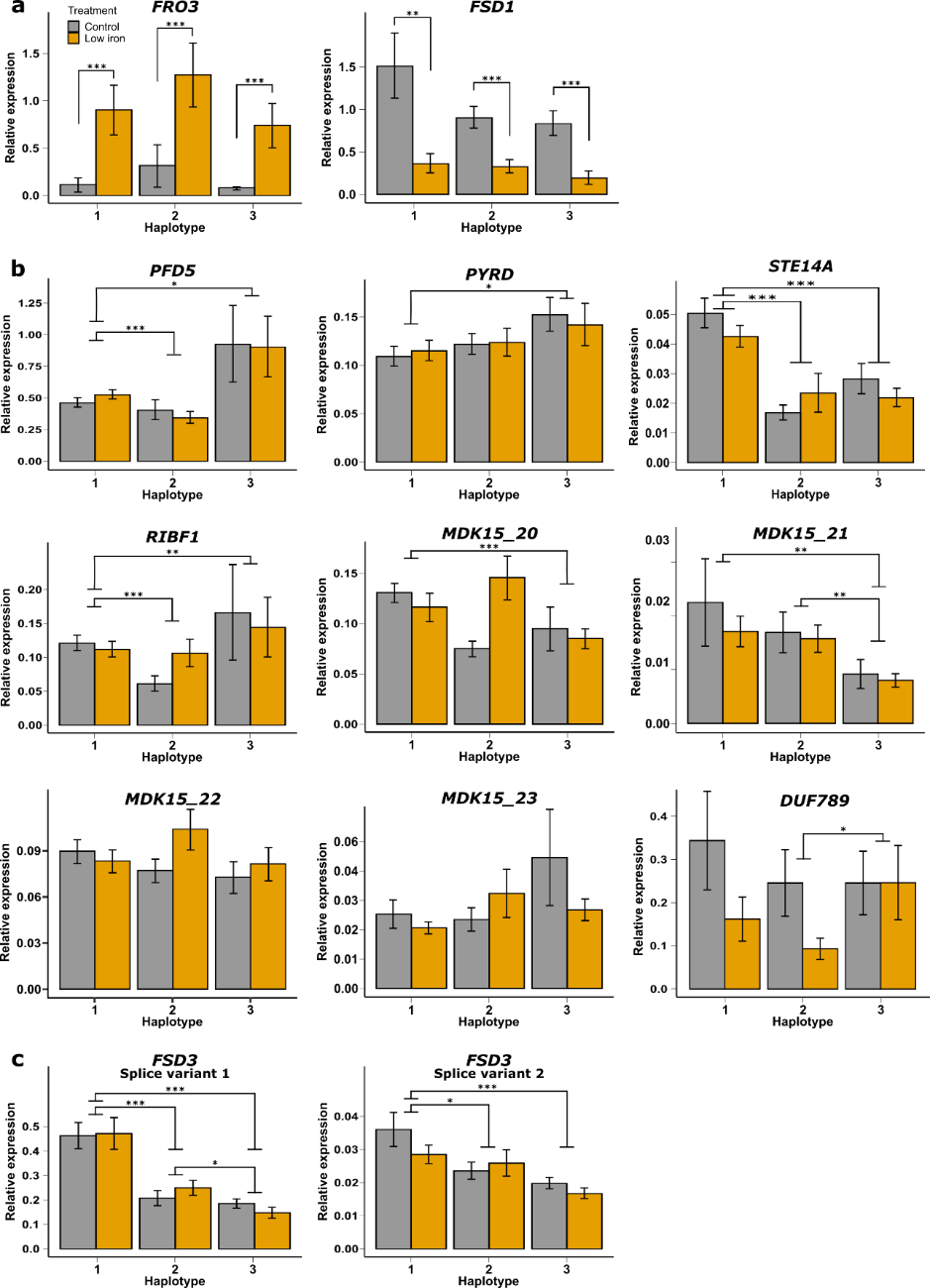

Figure S8: **Gene expression levels per haplotype group as measured by RT-qPCR.** The average gene expression levels at (21µm Fe^2+^ (Control, depicted in dark grey) and (1µm Fe^2+^) (Low iron, depicted in orange) per haplotype group for two iron deficiency marker genes (a), nine candidate genes from the GWAS (b) and two splice variants of *FSD3* (c). Average gene expression levels were taken from six natural accessions (with 5 replicates per accession per treatment) and normalized relative to three reference genes (*SAND*¸ *YLS8*, and *TIP41-like*). Significance of differences was assessed using two-sample t-tests on all Ct values from both treatments per haplotype. Error bars represent SE.**p <* 0*.*05; ***p <* 0*.*01; ****p <* 0*.*001.

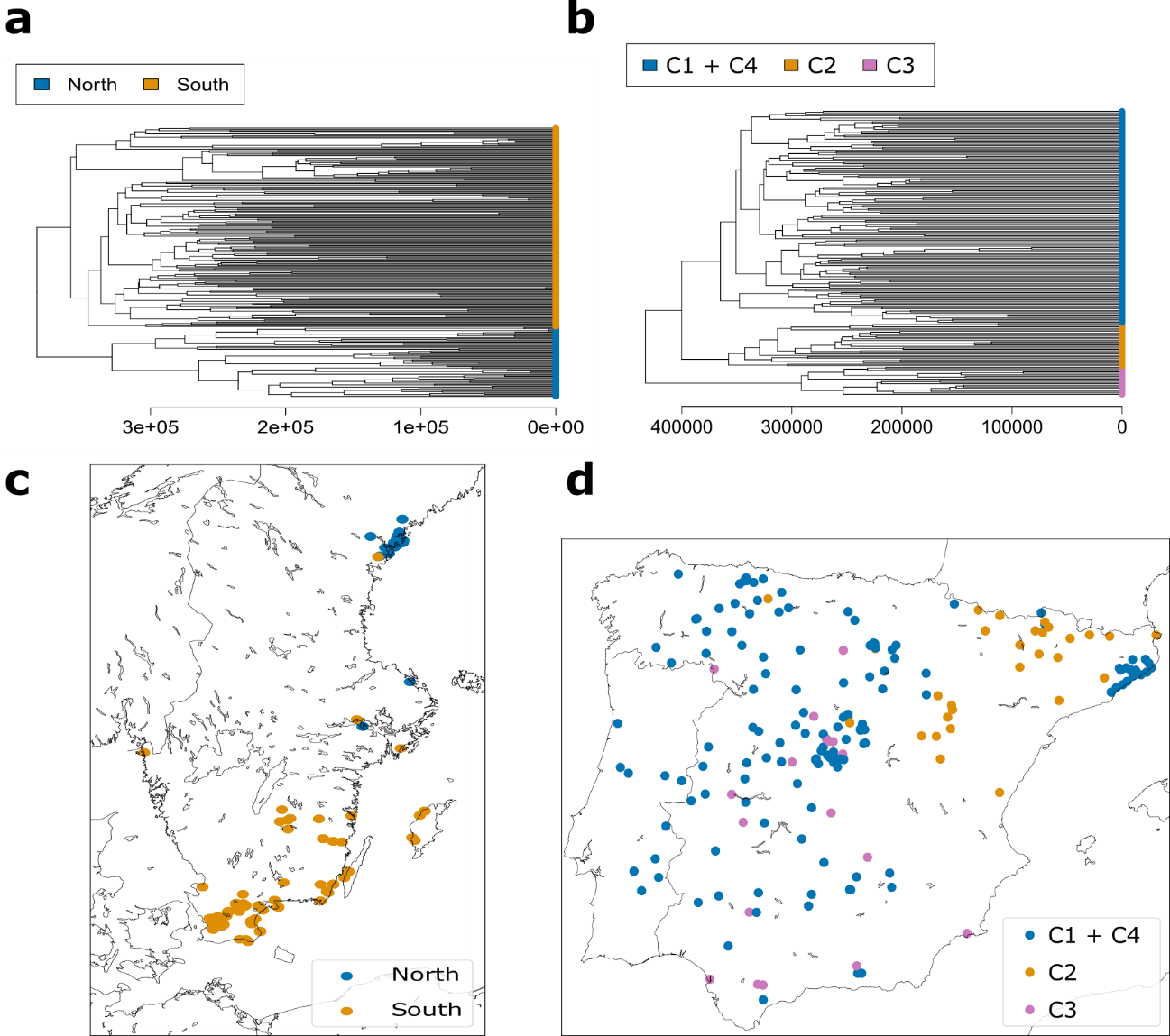

Figure S9: **Population structure of *A. thaliana* in Sweden and the Iberian peninsula.** (a) Dendrogram of Swedish accessions. (b) Dendrogram of Iberian accessions. Cluster names (C1, C2, C3 and C4) correspond to clusters previously reported^14^. (c) Geographical location of groups of Swedish accessions after cutting the dendrogram into two clusters. Groups correspond to accessions sampled in the north and south, as previously reported (Long et al., 2013). (d) Geographical location of groups of Iberian accessions after cutting the dendrogram into three clusters. Groups are distributed as previously reported (Tabas‐Madrid et al., 2018).

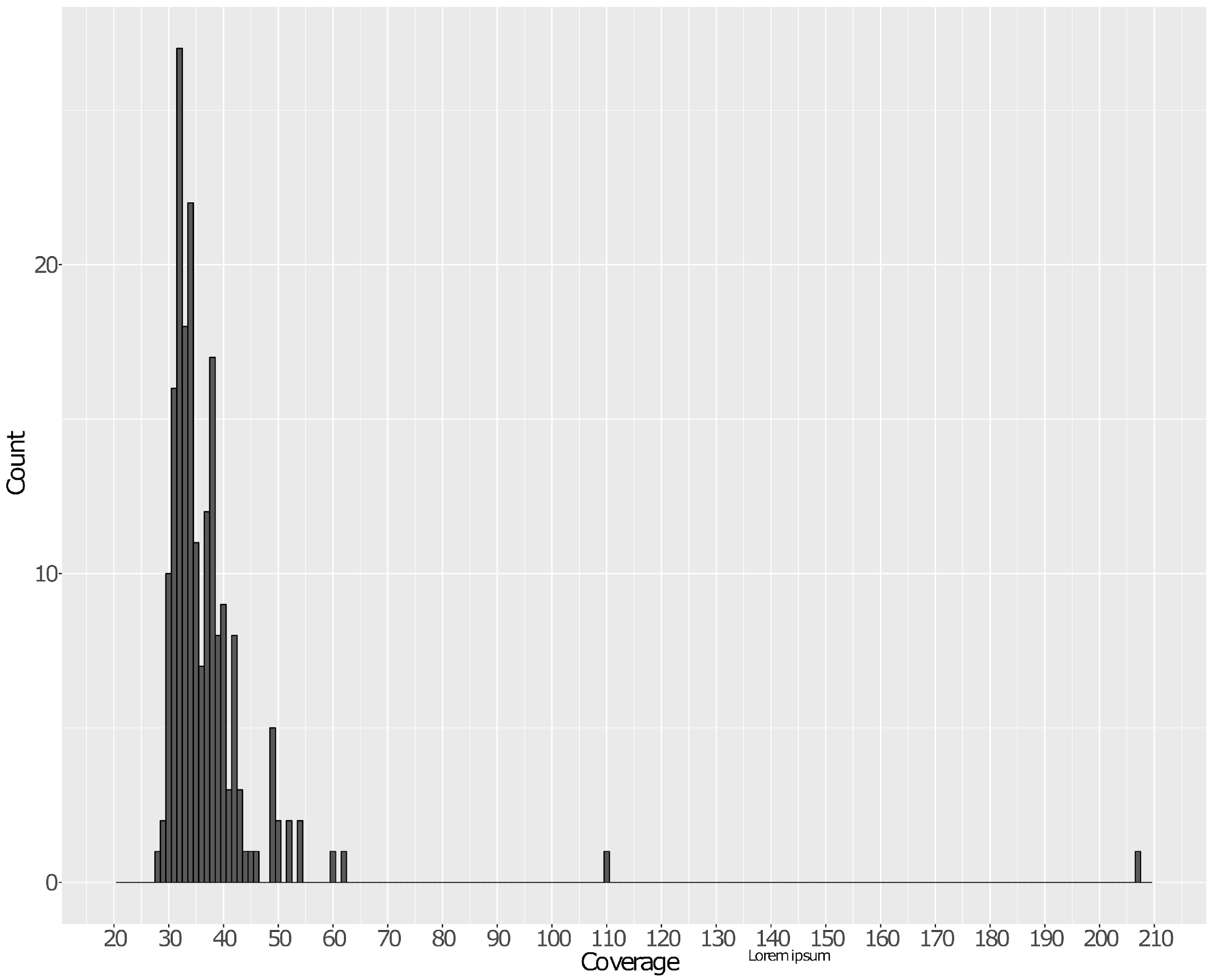

Figure S10: **Genomic coverage of each DartMap accession after pre-processing.** All samples have at least 28x coverage.

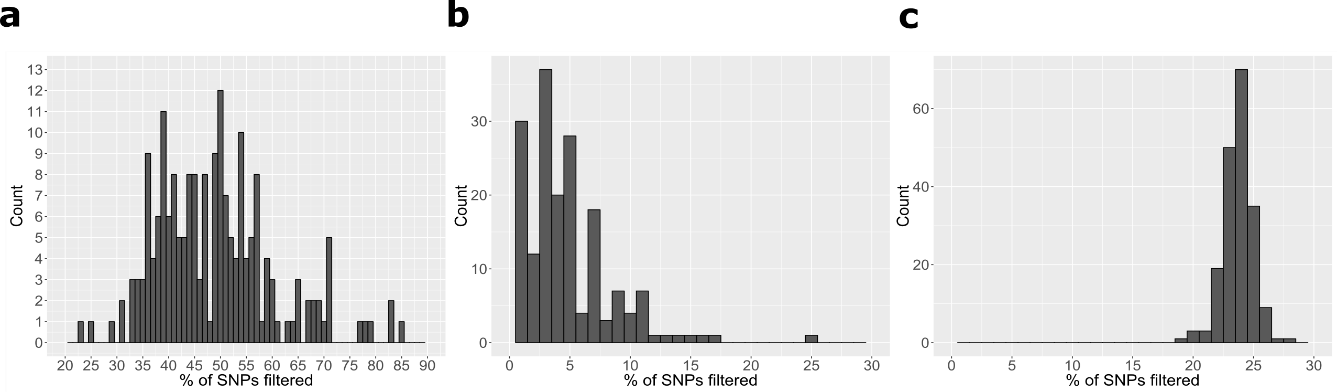

Figure S11: **Effect of filtering steps on number of retained SNPs for each sample.** The percentage of filtered SNPs is shown for the mitochondrial (a), chloroplast (b), and nuclear (c) callsets. The y-axes show the number of samples in which a particular percentage of SNPs (x-axes) were filtered.

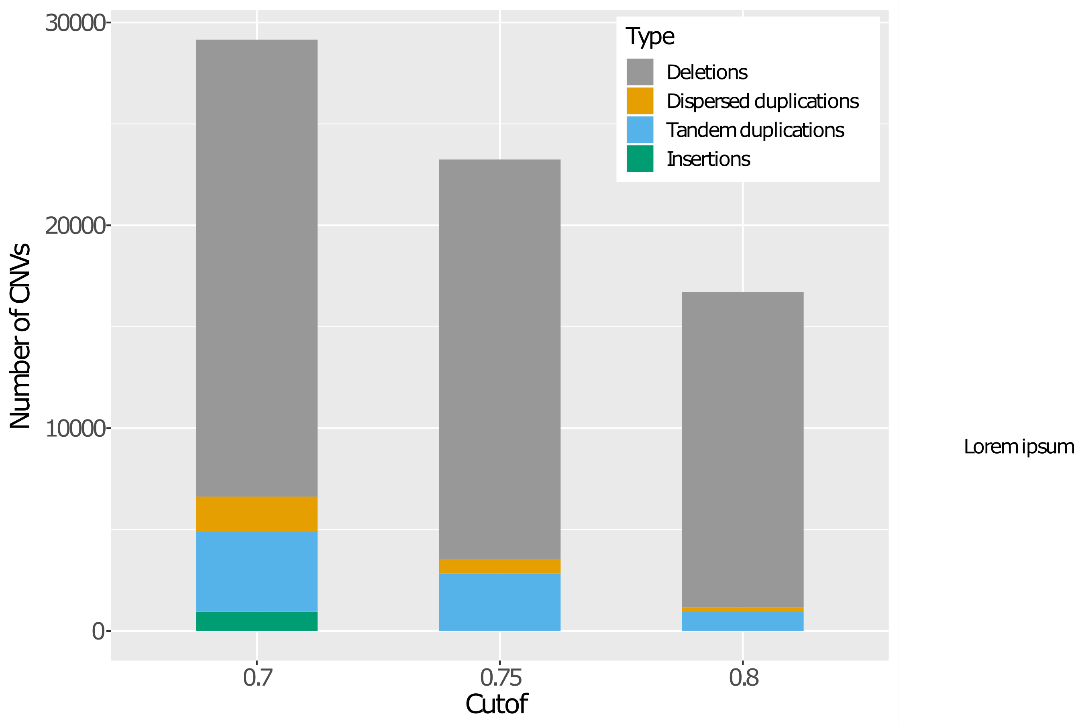

Figure S12: **Number and types of CNV detected at different cut-offs of Hecaton.** Insertions were not reported at high cut-offs, as Hecaton generally considers them to be inaccurate.

### Note S1: Supplementary Methods

Pre-processing genomic DNA sequence reads

To remove erroneous bases from the genomic DNA sequence data, we performed adapter and quality trimming using Cutadapt (Martin, 2011) (version 1.18). We clipped sequences that matched at least 90% of the total length of one of the adapter sequences provided in the NEBNext Multiplex Oligos for Illumina (Index Primers Set 1). In addition, we trimmed bases from the 5’ and 3’ ends of reads if they had a phred score of 20 or lower. Reads that were shorter than 100 bp after trimming were discarded.

Trimmed reads were aligned to a modified version of the A. thaliana Col-0 reference genome (TAIR10, European Nucleotide Accession number: GCA_000001735.2) which contained an improved assembly of the mitochondrial sequence (Sequence Read Archive accession number: BK010421) (Sloan et al., 2018) using bwa mem (Li, 2013) (version 0.7.10-r789) with default parameters. The resulting alignment files were sorted and indexed using samtools (Li et al., 2009) (version 1.3.1). We merged alignment files of libraries generated from the same accessions using Picard MarkDuplicates, which was called through the GATK suite (McKenna et al., 2010) (version 4.0.2.1). MarkDuplicates was also used to mark duplicate read pairs, using an optical duplicate pixel distance of 2500, as recommended in its documentation when working with patterned Illumina flowcells of the Hiseq 4000 platform.

We assessed whether each accession had adequate coverage to perform variant calling by computing the number of aligned bases of each accession using GATK CollectAlignmentSummaryMetrics. Each dataset had at least 28x coverage (Figure S10), which should be sufficient to accurately call SNPs, indels, and CNVs in a diploid plant species such as *A. thaliana*.

Filtering SNPs and indels

We filtered sets of SNPs and indels using two complementary approaches to remove likely false positive calls. First, we filtered the nuclear callset using GATK VariantRecalibrator and GATK ApplyVQSR (parameter –truth-sensitivity-filter-level set at 99.9). The set of variants called in a world-wide panel of 1135 *A. thaliana* accessions (The 1001 Genomes Consortium, 2016) were used as a training and truth set (prior=10.0) during recalibration. Second, we filtered variants using a set of hard thresholds as described in the documentation of GATK (https://gatk.broadinstitute.org/hc/en-us/articles/360035531112?id=6925). In order to choose appropriate thresholds, we generated density plots of six variant annotations known to be informative for identifying false positive calls (QD, FS, SOR, MQ, MQRankSum, and ReadPosRankSum) using a large set of SNPs and indels called in 1,506 A. thaliana samples of the 1,001 Genomes collection (The 1001 Genomes Consortium, 2016), an African collection (Durvasula et al., 2017), a Chinese collection (Zou et al., 2017), and a collection sampled from Madeira (Fulgione et al., 2018). To assess the effect of the filtering steps, we computed the percentage of SNP calls removed from each sample after filtering using Picard CollectVariantCallingMetrics (https://broadinstitute.github.io/picard/) (version 2.9.2). Most variants were retained after filtering (Figure S11), indicating that the majority of called SNPs are likely true positives.

Post-processing and validation of CNVs

CNV sites in each accession were genotyped by computing the read depth at the sites, relative to the rest of the chromosome they are located on, using duphold (Pedersen and Quinlan, 2019) (v0.1.1). Deletion sites were considered to be homozygous if they had a relative read depth lower than 0.25, heterozygous if they had a relative read depth between 0.25 and 0.75, and non-variant if they had a read depth higher than 0.75. Tandem and dispersed duplication sites were considered homozygous if they had a relative read depth higher than 1.75, heterozygous if they had a relative read depth between 1.25 and 1.75, and non-variant if they had a read depth below 1.25. As we cannot determine the genotype of insertions based on read depth, insertions were assigned a genotype of unknown/variant (./1) by default. After genotyping each site in each accession, we merged the call sets of each individual sample, merging CNVs that are likely to correspond to the same event. CNV calls were merged if they fulfilled the following conditions: they were of the same type; their breakpoints were located within 1000 bp of each other on both the 5’ and 3’ end; they share at least 50% reciprocal overlap with each other (does not apply to insertions); and the distance between the insertion sites is no more than 10 bp (only applies to dispersed duplications and insertions). The regions of the merged calls are defined as the union of the regions of the original calls. For instance, one call that covers positions 12-30 and a call that covers positions 14-32 are merged into a call covering positions 12-32. We excluded calls that were not supported by a change in read depth in any accession, as these are likely false positives.

Hecaton uses a random forest model to integrate the output of different callers and estimates a posterior probability estimate for CNVs, filtering those that fall below a certain threshold. To explore the effect of the filtering threshold, we generated sets using three different settings: 0.7, 0.75, and 0.8. No insertions were reported at thresholds of 0.75 and 0.8 (Figure S12), as these are difficult to accurately detect using short reads and therefore assigned a low probability. To include this type of CNV in our analysis, we chose to perform all further analyses using the set generated at a threshold of 0.7.

As sets of CNVs detected from short reads may contain spurious numbers of false positives and negatives (Cameron et al., 2019), we evaluated the sensitivity and precision of our detection approach by checking whether deletions reported in accessions were supported by a matching decrease in read depth. We were not able to use this method to assess the sensitivity and precision for duplications and insertions, as the expected increase in read depth at the affected regions can be caused by a number of other factors besides CNV, such as spurious alignments of reads in repetitive genomic regions. Read depth of regions predicted to be deleted in samples, relative to the 1000 bp flanks of such regions, was computed using duphold. Homozygous deletion calls were considered to be likely false positives if they had a relative read depth higher than 0.25, heterozygous deletions calls if they had a relative read depth lower than 0.25 or higher than 0.75, and non-variant calls if they had a read depth lower than 0.75. Note that the error rates computed using this method can be considered an upper bound, because reads of a sample may also fail to map to a region if it contains a large number of SNPs and indels in the sample genome relative to the reference (Schneeberger et al., 2009).

Investigating the effect of sequencing protocol on pairwise genetic distance

To investigate to what extent genetic distance between samples was influenced by differences in sequencing protocol, we modeled the genetic distance between different samples as a function of several technical co-variates. For each pair of samples *i* and *j*, we assume that the genetic distance *D_ij_* is linearly dependent on differences in sequencing coverage *C_ij_*, sequencing platform *P_ij_*, insert size *I_ij_*, and read length *R_ij_*:

$$D_{ij}=\beta_{0}+\beta_{1}C_{ij}+\beta_{2}P_{ij}+\beta_{3}I_{ij}+\beta_{4}R_{ij}$$

*β* parameters were estimated with the MRM function of the R ecodist package (Goslee and Urban, 2007) (v2.0.1), using genetic distances computed from the 1,506 *A. thaliana* samples used to define false positive SNP and indel calls. For the co-variate “sequencing platform”, we assigned a 1 to a sample if it was sequenced using Hiseq 3000 or a more recent platform, otherwise we assigned a 0. The reason for choosing Hiseq 3000 as a cut-off point is that Illumina platforms started using different flowcells (patterned instead of non-patterned) from that point on. We excluded insert size while modeling genetic distance based on SNPs, as not all samples were sequenced using paired-end libraries.
